# ParTIpy: A Scalable Framework for Archetypal Analysis and Pareto Task Inference

**DOI:** 10.1101/2025.09.08.674797

**Authors:** Philipp S. L. Schäfer, Leoni Zimmermann, Paul L. Burmedi, Avia Walfisch, Noa Goldenberg, Shira Yonassi, Einat Shaer Tamar, Miri Adler, Jovan Tanevski, Ricardo O. Ramirez Flores, Julio Saez-Rodriguez

**Affiliations:** Institute for Computational Biomedicine, Heidelberg University and Heidelberg University Hospital, Heidelberg, Germany; European Molecular Biology Laboratory, European Bioinformatics Institute (EMBL-EBI), Hinxton, Cambridgeshire, U.K; Department of Genetics, Silberman Institute of Life Sciences; and Department of Immunology and Cancer Research, Faculty of Medicine, The Hebrew University of Jerusalem, Jerusalem, Israel; Department of Biological Chemistry, Silberman Institute of Life Sciences, The Hebrew University of Jerusalem, Jerusalem, Israel

**Author notes:** These authors jointly supervised the work (Ricardo O. Ramirez Flores, Julio Saez-Rodriguez).

**Keywords:** Archetypal Analysis, Pareto Task Inference, Single-Cell Transcriptomics, Spatial Transcriptomics, Trade-Offs, Resource Allocation, Division of Labor

## Abstract

**Motivation:** Trade-offs between different functions or tasks are pervasive across scales in biological systems. For example, individual cells cannot perform all possible functions simultaneously; instead they allocate limited resources to specialize in subsets of tasks by activating specific gene expression programs. Pareto Task Inference (ParTI) is a framework for analyzing biological trade-offs grounded in the theory of multi-objective optimality. However, existing software implementations of ParTI lack scalability to large datasets and do not integrate well with standard biological data analysis workflows, especially in the context of single-cell transcriptomics, limiting broader adoption.

**Results:** We have developed ParTIpy (Pareto Task Inference in Python), an open-source Python package that combines advances in optimization and coreset methods to scale archetypal analysis, the primary algorithm underlying ParTI, to large-scale datasets. By providing additional tools to characterize archetypes, comprehensive documentation, and adopting standard scverse data structures, ParTIpy facilitates seamless integration into existing analysis workflows and broadens accessibility, particularly within the single-cell community. We demonstrate how ParTIpy can be used to study intra-cell-type gene expression variability through the lens of task allocation, offering a principled alternative to methods that impose discrete cell state classifications on inherently continuous variation.

**Availability and implementation:** ParTIpy’s open-source code is available on GitHub (https://github.com/saezlab/ParTIpy) and pypi (https://pypi.org/project/partipy). Documentation is available at https://partipy.readthedocs.io. The code to reproduce the results of this paper is on GitHub (https://github.com/saezlab/ParTIpy_paper)

## 1. Introduction

Trade-offs between different functions or tasks are pervasive across scales in biological systems. At the morphological level, the beaks of Darwin’s finches are a classical example: Their beaks cannot be optimal for both cracking seeds and picking insects, leading to specialization or compromise across species (*1*, *2*). At the cellular scale, individual cells cannot perform all functions simultaneously; instead they allocate limited resources to specialize in subsets of tasks (*3–7*). Pareto Task Inference (ParTI) is a framework for analyzing trade-offs grounded in the theory of multi-objective optimality (*2*, *3*). It assumes that evolution drives phenotypes toward Pareto optimality, where improvement in one task necessarily worsens performance in another. Under the assumption of Pareto optimality^1^, phenotypes (e.g. organisms, proteins, or cells), described by traits with trade-offs (e.g. beaks of finches, kinetic properties, or gene expression), lie within a polytope^2^ that has as many vertices as tasks being optimized (*2*). Leveraging this geometric structure, ParTI infers the underlying tasks by fitting a polytope to phenotypic measurements, where each vertex, referred to as an archetype, represents a phenotype optimized for a specific task. For example, in the context of single-cell transcriptomics, ParTI fits a polytope enclosing cells in gene expression space, with vertices (archetypes) representing cells optimized for distinct tasks. The gene expression profile of each archetype can then be analyzed to infer the specific tasks. The polytopes can be efficiently inferred using archetypal analysis, a dimensionality reduction technique that identifies extreme points of a dataset, corresponding to the archetypes, and models the dataset as non-negative linear combinations of these archetypes (*8–10*).

Numerous studies have demonstrated the utility of archetypal analysis for modeling biological systems, particularly at the single-cell level (*4–7*, *11–16*), revealing trade-offs in cellular systems such as hepatocytes (*5*), fibroblasts (*6*), and cancer cells (*16*). In addition, archetypal analysis has been applied to spatial transcriptomics and proteomics data, where instead of focusing on individual cells, the objective is to identify groups of spatially co-localized cells that cooperate to perform specific tasks, thus revealing a division of labor at the niche level (*17–20*).

To enable application of ParTI in biology, the `ParTI` MATLAB package (*3*) was developed, combining algorithms for polytope inference with tools for archetype characterization into a principled workflow for Pareto task inference. However, the software does not scale to large data sets, in particular those generated with high-throughput single-cell technologies. Moreover, it requires a commercial license (MATLAB), hindering broader adoption.

To address these limitations, we developed ParTIpy (Pareto Task Inference in Python), an open-source Python package that combines algorithmic innovations in initialization and optimization with the use of coresets (*9*, *21–23*) to scale archetypal analysis to large-scale datasets. Although ParTIpy is broadly applicable, we focus here on its integration with single-cell analysis workflows in the scverse (*24*). In particular, we show how ParTIpy can be used to study intra-cell-type variability through the lens of optimal resource allocation and division of labor, offering a principled alternative to approaches that impose discrete cell state classifications onto inherently continuous variation.

## 2. Implementation & Results

ParTIpy provides a modular framework for applying archetypal analysis to multivariate data, with various options for initialization and optimization, and support for relaxing model constraints. It supports the use of coresets to scale analysis to large datasets and includes metrics to help determine the optimal number of archetypes. To support single-cell transcriptomic applications, ParTIpy integrates downstream functionalities for interpreting archetypes, including functional enrichment, cross-condition comparisons, spatial mapping, and inference of archetypal crosstalk networks via ligand-receptor co-expression analysis (Figure 1).

**Figure 1.**
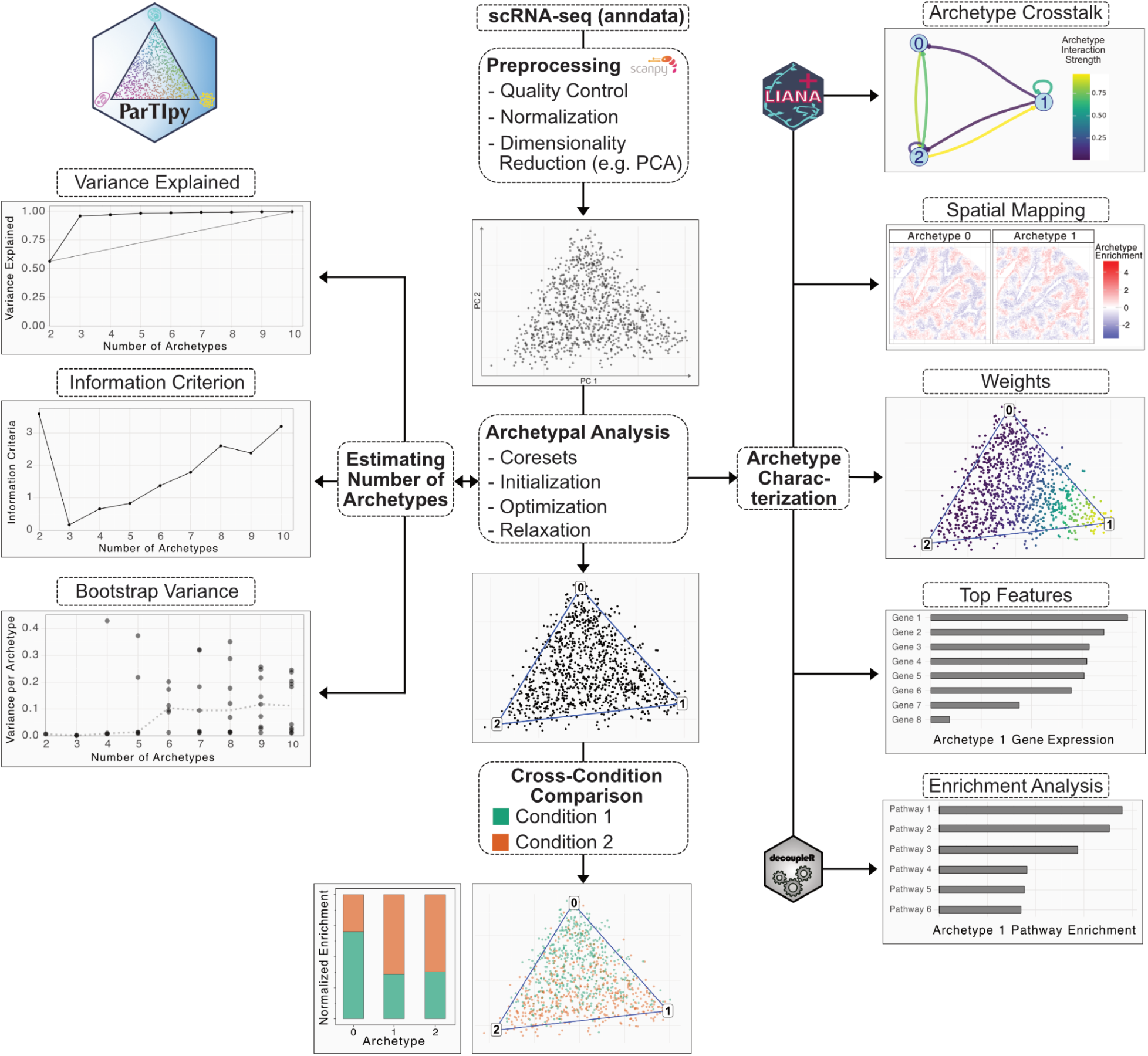
Overview of the ParTIpy framework.

### 2.1 Archetypal Analysis

Archetypal analysis finds K extreme points within the data distribution (convex hull of the data) called archetypes, such that all other data points can be described as mixtures (convex combinations) of these archetypes (*8–10*). Formally, the archetypes *Z*∈*R* ^*K* ×*D*^ are defined as convex combinations of the data points, *Z* = *BX* where *X*∈*R*^*N* ×*D*^ is the data matrix with N cells, D features and *B*∈*R*^*K*×*N*^ is a non-negative weight matrix where each row sums to one (row stochastic). This constraint ensures that the archetypes lie within the convex hull of the dataset *X*, forming *K* vertices of a polytope that approximates the data distribution. In the context of ParTI, each vertex corresponds to an inferred archetype that represents a distinct task a cell may be specialized in, reflecting an underlying division of labor. Each data point is then reconstructed as a convex combination of the archetypes, *X*≈*A Z* where ^*N*×*K*^ is also row-stochastic. Putting everything together, archetypal analysis seeks a matrix *B* that defines *K* archetypes as extreme points within the data distribution, and a matrix *A* that reconstructs each data point as a convex combination of these *K* archetypes (see Equation 1).

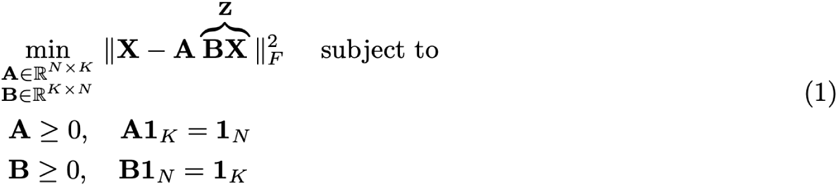

However, when only a few cells have been sampled near the boundaries of the true phenotypic space, the convex hull of the observed data may underestimate the true range of variation. Such limited sampling near the boundaries can arise, for example, when the sample size is small or when the population is dominated by generalist cells rather than cells specialized in tasks. In these cases, relaxing the convexity constraint can help recover the true archetypes. This is achieved by introducing a relaxation parameter δ (see Equation 2) (*9*), where larger values of δ permit archetypes to lie farther from the data cloud (see Supplementary Figure S1).

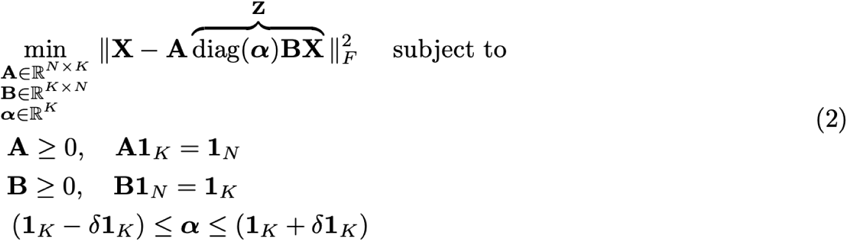

The non-convexity of the objectives in Equations 1 and 2 gives rise to challenging optimization landscapes, where solution quality is highly sensitive to initialization. To address this, a range of initialization and optimization strategies have been developed to improve convergence and reconstruction accuracy (*10*). We have implemented and benchmarked three different initialization algorithms and two different optimization algorithms (see Supplementary Methods Sections 2.3 and 2.4). Using 21 single-cell transcriptome datasets, each corresponding to a single cell type, from three independent studies (Supplementary Table S1), we tested which combination of initialization and optimization algorithm offers the best trade-off between reconstruction accuracy and runtime (see Supplementary Methods Section 5.1). We found that using the principal convex hull analysis (PCHA) optimization algorithm (*9*) and the Archetypal++ (AA++) initialization (*23*) yields the best results (Figure 2A, Supplementary Figure S2). However, if runtime is a limiting factor, the more efficient Frank-Wolfe (FW) algorithm (*21*) provides a practical alternative.

**Figure 2.**
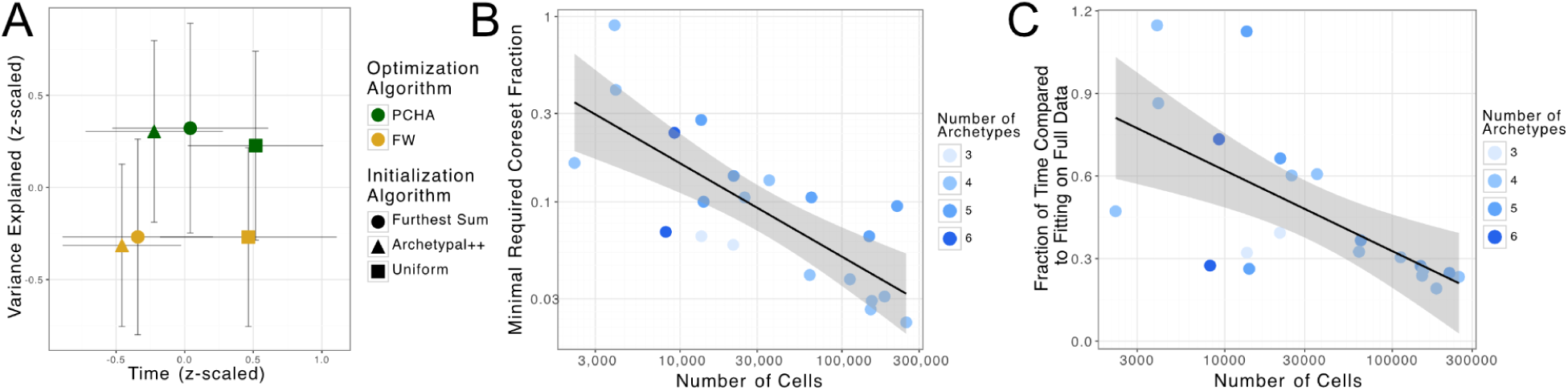
Benchmarking optimization parameters and coreset settings. (A) Summary of benchmark results for initialization and optimization algorithms across all datasets and cell types shown in Table 1. (B) Minimal coreset size required to capture at least 95% of the variance explained by the full-data model. (C) Fraction of run time for optimization with the minimal required coreset size compared to the full-data model.

To further improve the scalability of ParTIpy, we implemented an algorithm for constructing coresets specifically designed for archetypal analysis (see Supplementary Methods Section 2.5) (*22*). A coreset is a weighted subset of the data such that optimizing the objective on the coreset yields results comparable to those obtained using the full dataset. This requires incorporating a diagonal weight matrix *W* into the objective function, where *X̂* denotes the subsampled data matrix representing the coreset (Equation 3).

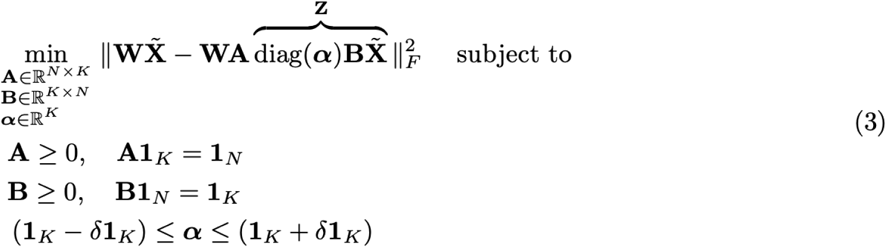

To support this formulation, we modified the optimization algorithms accordingly (see Supplementary Methods Section 2.5). Using the same datasets employed for benchmarking initialization and optimization strategies (see Supplementary Table S1), we evaluated the minimum coreset size required to achieve comparable results to full-data optimization under realistic conditions, defined as the smallest coreset capturing at least 95% of the variance explained by the full-data model. As shown in Figure 2B-C and Supplementary Figure S4, we found that the required coreset fraction decreases with increasing dataset size. For instance, when analyzing datasets with over 100,000 cells, using only 1–10% of the data was sufficient on average, resulting in approximately a fourfold reduction in runtime.

To validate that ParTIpy produces results comparable to the original `ParTI` MATLAB package (*3*), we analyzed three distinct single-cell datasets (see Supplementary Table S2) and compared the resulting archetypes. As shown in Supplementary Figure S3, we observed a high degree of concordance between the two implementations.

Altogether, ParTIpy offers a flexible and scalable implementation of archetypal analysis that supports various initialization and optimization schemes, includes convexity relaxation, and introduces coreset-based acceleration, extending the original ParTI framework while preserving result concordance.

### 2.2 Choosing the Number of Archetypes

To determine the appropriate number of archetypes, we implemented three complementary criteria, each evaluated over a range of candidate values (see Figure 1, left panel and Supplementary Methods Section 3).

First, we compute the variance explained. An elbow in this curve suggests that adding additional archetypes yields diminishing returns. Second, we use an information-theoretic criterion (*25*) where lower values indicate a favorable trade-off between model complexity (i.e., the number of archetypes) and variance explained. Third, we assess archetype robustness via bootstrapping: we apply archetypal analysis to multiple bootstrap samples, align the resulting archetypes across replicates, and compute the positional variance of each archetype (*3*, *4*). A lower mean variance indicates greater robustness.

### 2.3 Characterizing the Archetypes

To characterize the archetypes, we first compute the distance of each cell to each archetype. Then we transform these distances into weights using a squared exponential kernel (see Figure 3D and Supplementary Figure S5D). By multiplying this weight matrix with the feature matrix, we obtain one aggregated feature profile per archetype (see Supplementary Methods Section 4).

**Figure 3.**
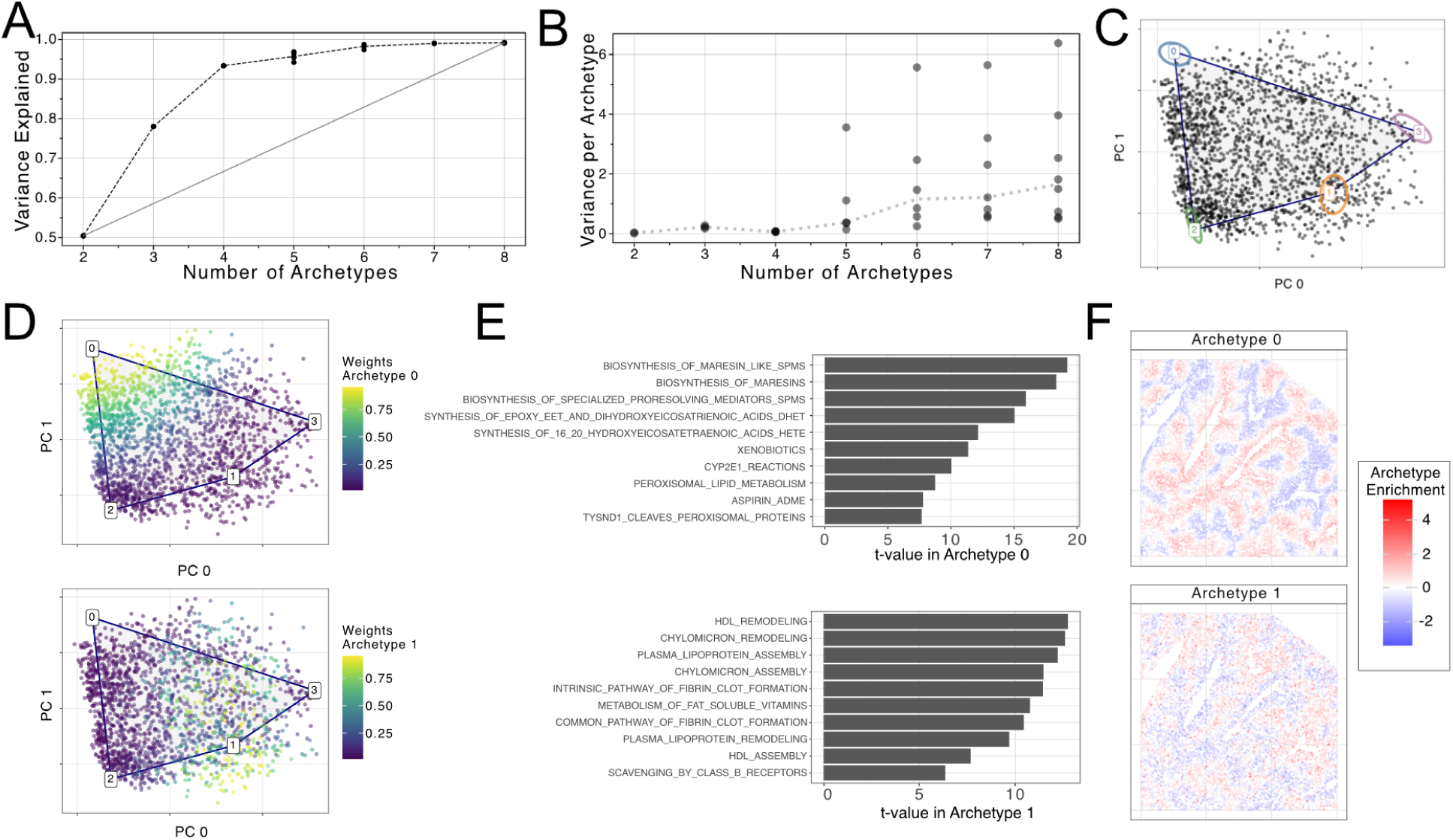
Example analysis on hepatocyte data from (29).(A) Variance explained plotted for two to nine archetypes. (B) Bootstrap variance plotted for two to nine archetypes. (C) Two-dimensional PC plot showing single-cells in black, and the archetype in different colors; contours represent a 95% interval derived from the bootstrap samples. (D) Two-dimensional PC plot showing the weights used to characterize archetypes 0 and 1. (E) Barplots showing the top enriched Reactome pathways for archetypes 0 and 1. (F) Spatial enrichment of archetype 0 and 1.

For single-cell transcriptomic applications, archetypes can be characterized by directly inspecting the aggregated expression profiles or by performing pathway enrichment analysis. To facilitate the latter, ParTIpy seamlessly integrates with functions from decoupler-py (*26*), enabling pathway enrichment analysis within the framework.

Having identified distinct archetypes and their associated biological functions, a natural next step is to ask what gives rise to this observed division of labor. One plausible explanation is the existence of spatial (chemical) gradients that render certain tasks more favorable in specific tissue regions (*5*). When spatial transcriptomic data from the same biological context are available, this hypothesis can be tested by mapping archetypes onto the spatial dataset, as described in Supplementary Methods Section 4.2. Another potential driver of functional specialization is cell-cell communication, such as lateral inhibition, where a cell performing a given task inhibits nearby cells from adopting the same task (*6*). To investigate whether such interactions underlie the observed division of labor, we infer potential signaling between archetypes by integrating their characteristic expression profiles with ligand–receptor databases (*27*, *28*), following the workflow adapted from (*6*) (see Supplementary Methods Section 4.3).

### 2.4 Using ParTIpy to Characterize Hepatocytes

To illustrate the use of ParTIpy, we applied it to a single-cell RNA-seq dataset of hepatocytes sampled along the porto-central axis of the liver lobule (see Supplementary Table S2) (*29*). Hepatocytes exhibit spatially constrained functional trade-offs, balancing distinct metabolic, biosynthetic, and detoxification processes (*30*), and ParTI has previously been used to infer archetypes related to this division of labor (*5*). We first used the three archetype selection criteria introduced above to determine the number of archetypes (Figure 3A-B, Supplementary Figure S5A). Based on these metrics, we selected four archetypes, as increasing beyond four yielded negligible gains in variance explained and led to reduced robustness. To characterize the resulting archetypes, we performed pathway enrichment analysis using decoupler-py (*26*) and the Reactome database (*31*). Archetype 0, for example, was associated with drug and xenobiotic metabolism, while archetype 1 was enriched for lipid metabolism (Figure 3E, Supplementary Figure S4J-M). To investigate potential mechanisms underlying this division of labor, we mapped the archetypes onto single-cell resolved spatial transcriptomics data (Vizgen MERFISH Mouse Liver Map January 2022). This analysis revealed that archetype 0 is localized near the central vein, whereas archetype 1 is enriched near the portal node (see Figure 3F), consistent with previous hepatocyte archetype analysis findings (*5*)and established zonation patterns (*30*). These results suggest that the observed functional specialization is driven by spatial gradients of oxygen and other metabolites along the porto-central axis.

## 3. Discussion

Here we present ParTIpy, an open-source Python package that scales archetypal analysis by implementing state-of-the-art initialization and optimization techniques in combination with coresets, enabling the application of ParTI to large datasets, for example, the ones generated by modern single-cell transcriptomics technologies (*32*). Applied to single-cell data, ParTIpy reveals how individual cells distribute across competing transcriptional programs, offering a complementary perspective to conventional clustering approaches. While we mainly presented examples on dissociated single-cell data, ParTIpy is broadly applicable to other biological contexts; for example, our online tutorials demonstrate its use with spatial transcriptomics.

ParTIpy implements two optimization algorithms for archetypal analysis (*10*), selected from the broader collection for their flexibility and efficiency, and our benchmarks and applications confirm that they provide the scalability required for modern single-cell datasets. Our gradient-based optimization algorithms readily accommodate modifications to the objective function, including the convexity relaxation that allows archetypes to extend beyond the data cloud. One notable extension of archetypal analysis is deep archetypal analysis (*33–36*), in which an encoder–decoder neural network is jointly learned with archetypes defined in the latent space of the encoder. This approach is motivated by its ability to capture non-linear structure in the data and its scalability due to mini-batch gradient descent. Rather than relying on non-linear models, ParTIpy achieves scalability through coreset construction and adopts linear dimensionality reduction, which ensures that the archetypes correspond to extremal points in the original trait space, consistent with the theoretical foundations of ParTI.

In conclusion, ParTIpy combines computational efficiency with biological interpretability in a versatile framework for exploring functional trade-offs in high-dimensional biological data. With dedicated tools for archetype characterization, seamless integration into Python-based workflows, and comprehensive documentation (https://partipy.readthedocs.io), it broadens access to ParTI for the research community.

## 4. Acknowledgements

P.S.L.S. has received funding from the Deutsche Forschungsgemeinschaft under grant agreement SPP 2395.

We thank Katharina Mikulik and Bárbara Zita Peters Couto for valuable discussions and constructive feedback on the manuscript.

## 5. Conflict of interests

JSR reports in the last 3 years funding from GSK and Pfizer and fees/honoraria from Travere Therapeutics, Stadapharm, Astex, Owkin, Pfizer, Grunenthal, Tempus and Moderna.

## 6. Authors contributions

P.S.L.S. and R.O.R.F. conceived the idea. P.S.L.S. and L.Z. wrote and tested the software package. P.S.L.S., L.Z, and P.L.B. worked on the benchmark and application with support from J.T. and R.O.R.F. P.S.L.S. and R.O.R.F. wrote the manuscript. A.W., N.G., S.Y., E.S.T., and M.A. compared ParTIpy to the ParTI MATLAB package. R.O.R.F., and J.S.R. supervised the work. J.S.R. acquired funds.

## 7. Data availability

For the datasets used in the benchmarking part, see Supplementary Table S1. For the datasets used to compare the ParTI MATLAB package and ParTIpy, see Supplementary Table S2. The scripts used to download the data are here https://github.com/saezlab/ParTIpy_paper/blob/main/code/utils/data_utils.py.

## 9. Supplementary Figures

**Supplementary Figure S1.**
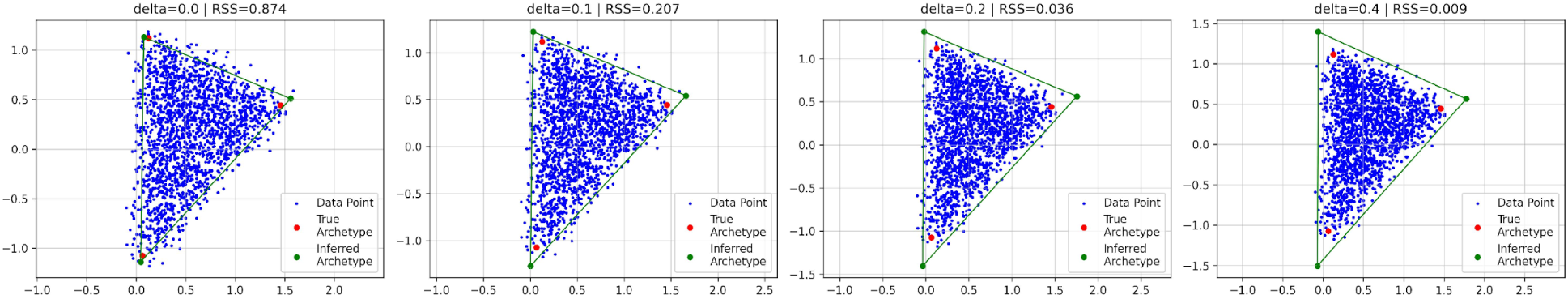
Effect the δ parameter. As δ increases, archetypes deviate further from the true *positions, while the residual sum of squares (RSS) decreases toward zero*.

**Supplementary Figure S2.**
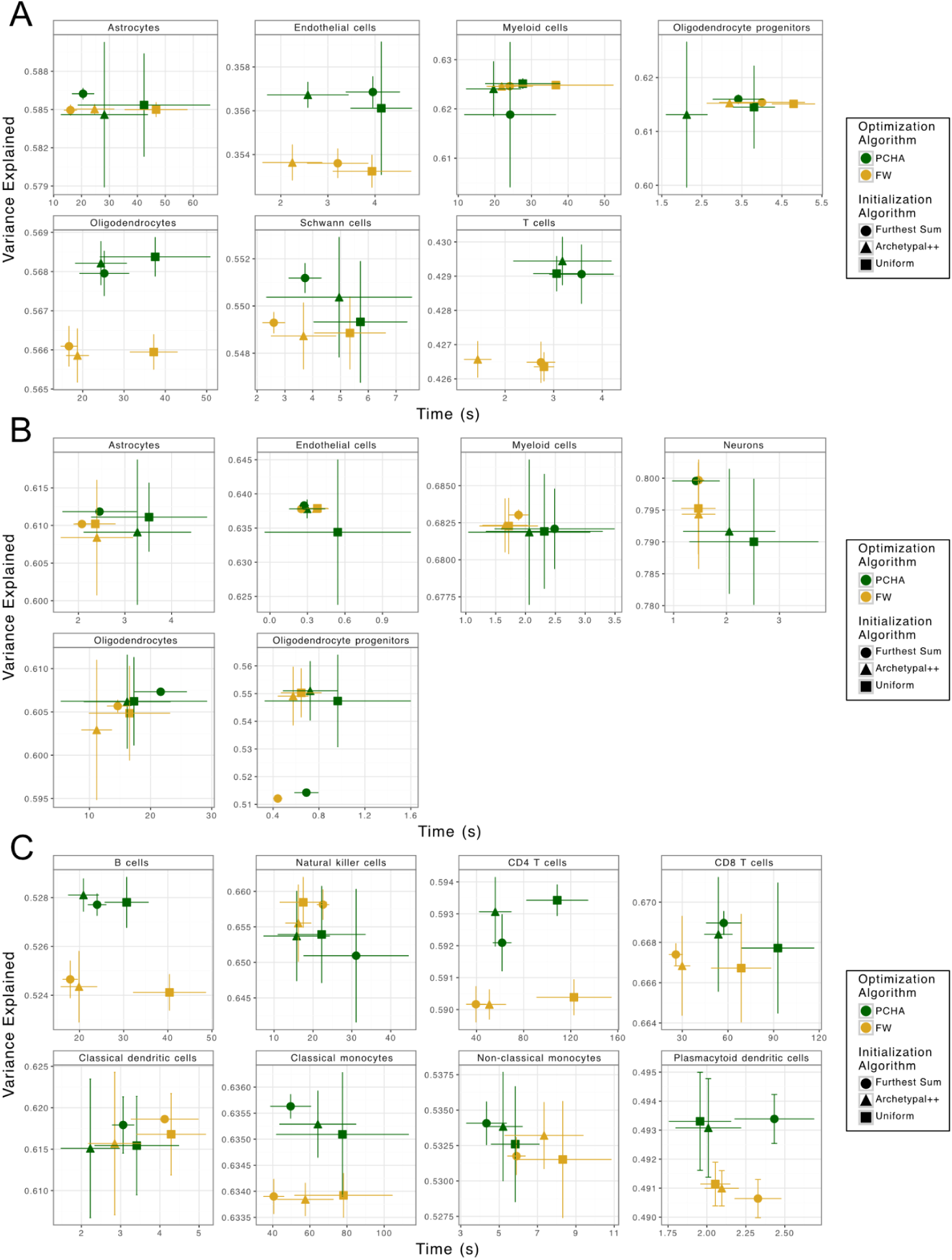
Benchmarking initialization and optimization algorithms for archetypal analysis. Results per study: A) Kukanja et al. 2024, (B) Lerma-Martin et al. 2024, (C) Perez et al. 2022. Each point denotes the mean runtime and mean variance explained across 20 runs with different random seeds; error bars indicate standard deviations across those same runs.

**Supplementary Figure S3.**
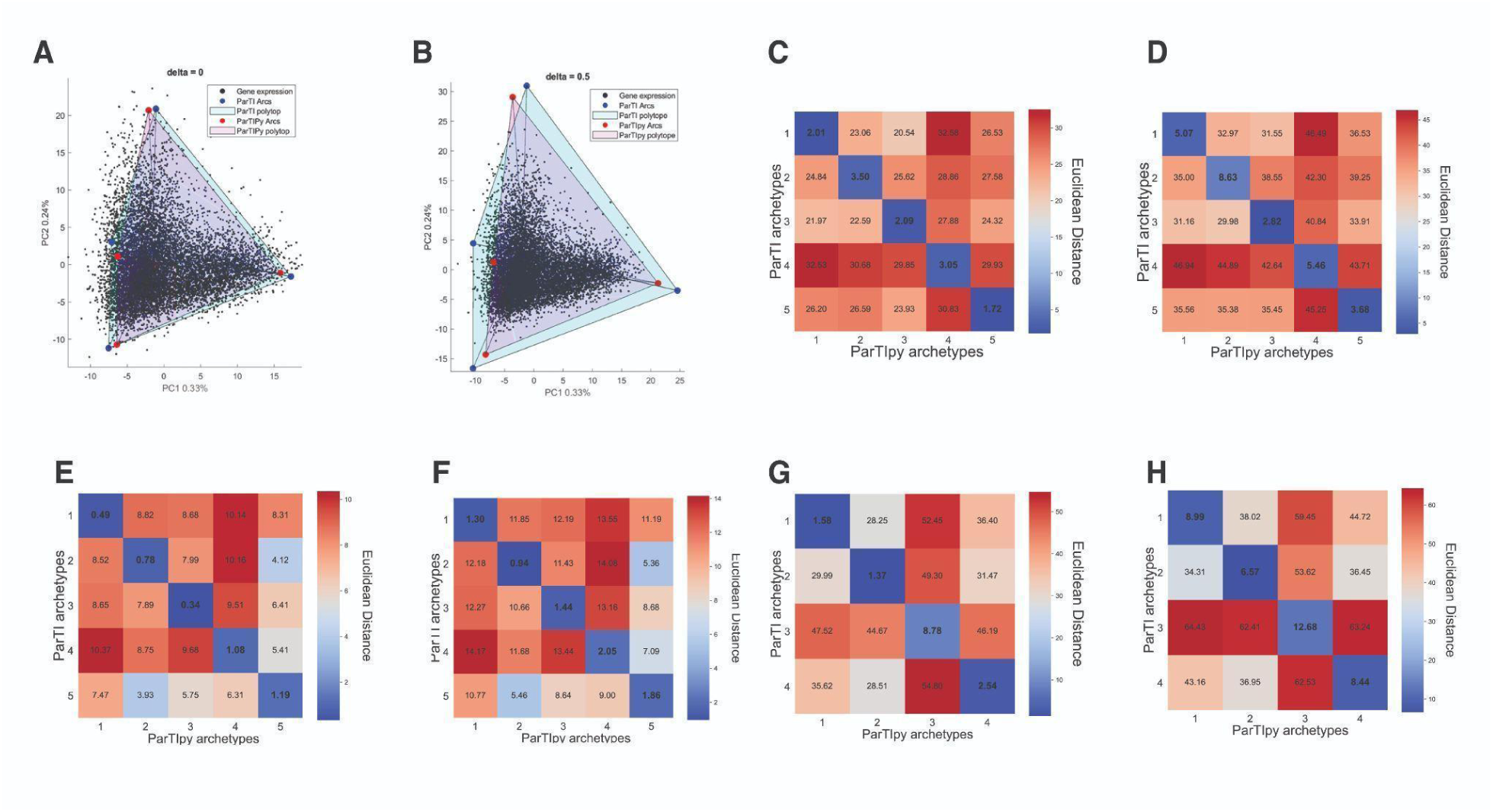
Comparison of `ParTI` MATLAB package and ParTIpy. (A–D) Heart fibroblasts: snRNA-seq profiles from hypertrophic cardiomyopathy patients (37). ParTIpy simplex (purple) overlaps the ParTI simplex (green) at δ = 0. 0 (A) and δ = 0. 5 (B). (C-D) Minimal Euclidean-distance heat-maps show only small gaps between the corresponding archetypes for δ = 0. 0 (C) and δ = 0. 5 (D). (E–F) Spinal-cord myeloid cells: 10× Xenium spatial-transcriptomics data from multiple-sclerosis patient spinal cords (38). Minimal Euclidean-distance heat-maps reveal small gaps between corresponding archetypes. (G–H) scRNA-seq of hepatocytes (29) likewise shows close agreement between methods.

**Supplementary Figure S4.**
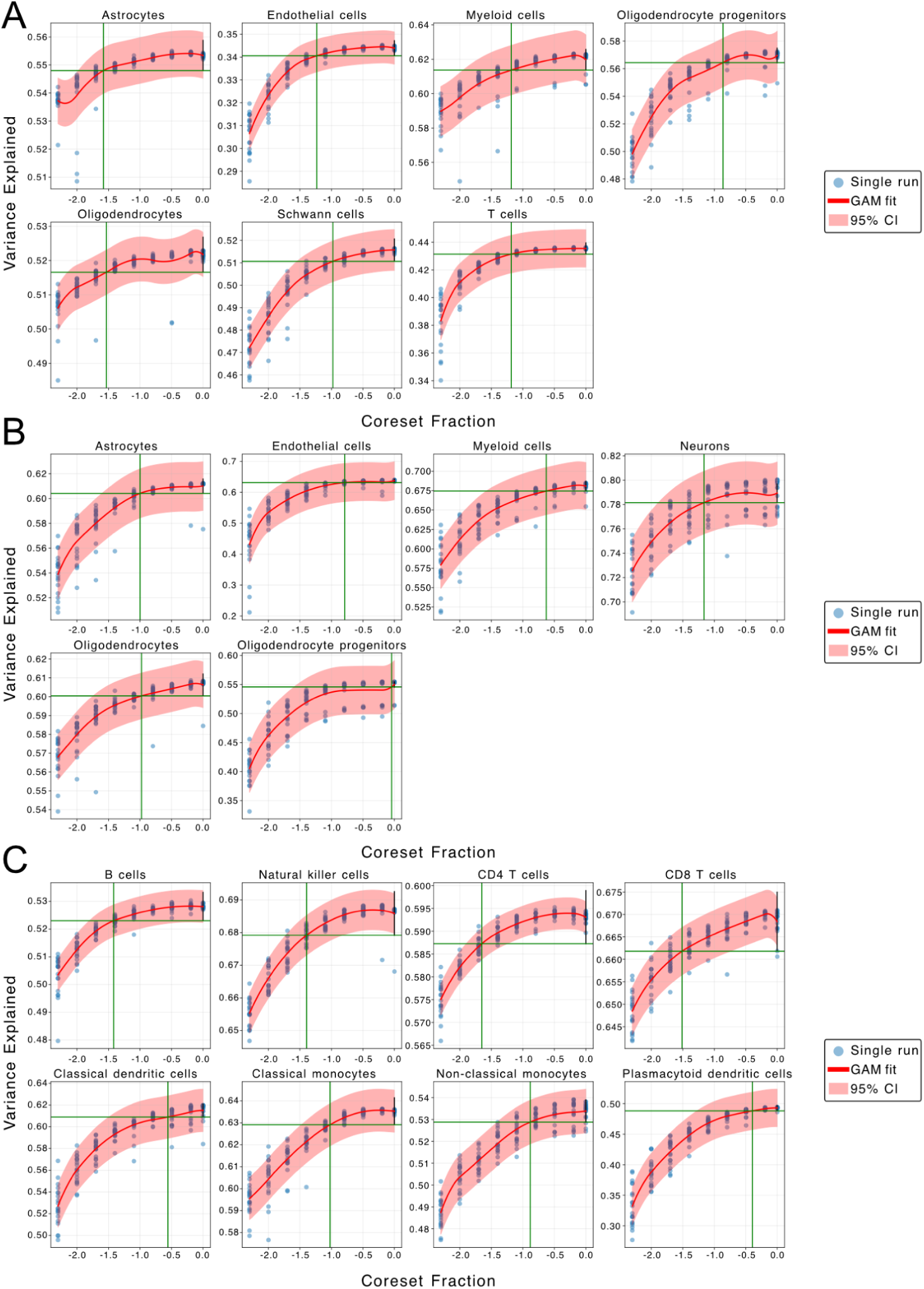
Evaluating the performance of coresets. Results per study: Kukanja et al. 2024 (38) (A), Lerma-Martin et al. 2024 (39) (B), Perez et al. 2022 (40) (C). We tested coreset sizes of 1.0, 0.64, 0.32, 0.16, 0.08, 0.04, 0.02, 0.01, and 0.005. For each combination of coreset size and cell type, optimization was repeated 20 times with distinct random seeds. The red line indicates a fit obtained using a generalized additive model (GAM). The green line indicates the smallest coreset size required to capture 95% of the variance explained by the full-data model.

**Supplementary Figure S5.**
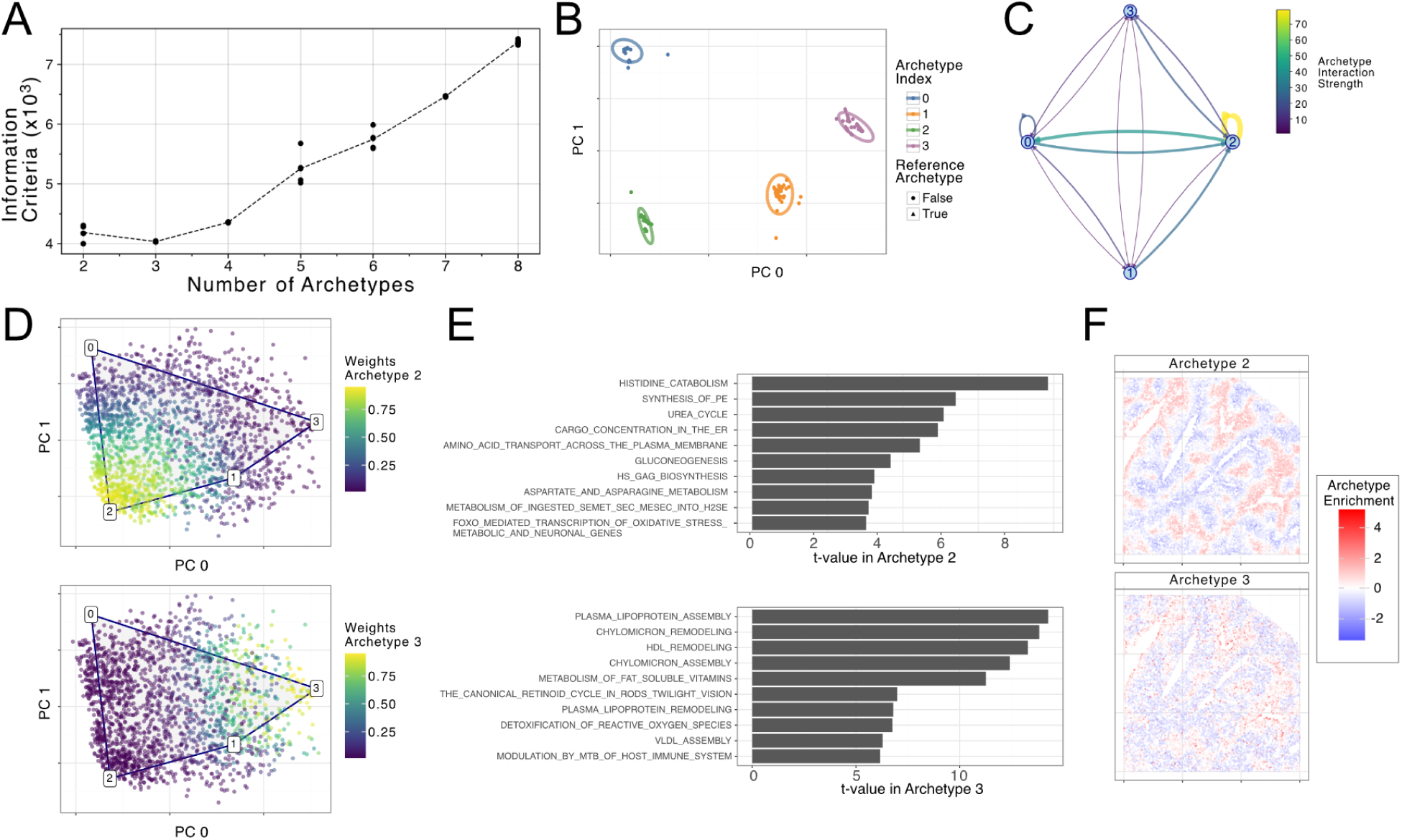
Example analysis on hepatocyte data from (29). (A) Information criterion for archetypal analysis plotted for two to nine archetypes. (B) Two-dimensional PC plot showing the archetypal positions obtained from performing the optimization on 30 different bootstrap samples. (C) Crosstalk analysis shows strong self-interaction in archetype 2, and strong interaction between archetypes 0 and 2. (D) Two-dimensional PC plots showing the weights used to characterize archetypes 2 and 3. (E) Barplots showing the top enriched Reactome pathways for archetypes 2 and 3. (N) Spatial mapping of the archetypes 2 and 3.

## 10. Supplementary Tables

**Supplementary Table S1.**
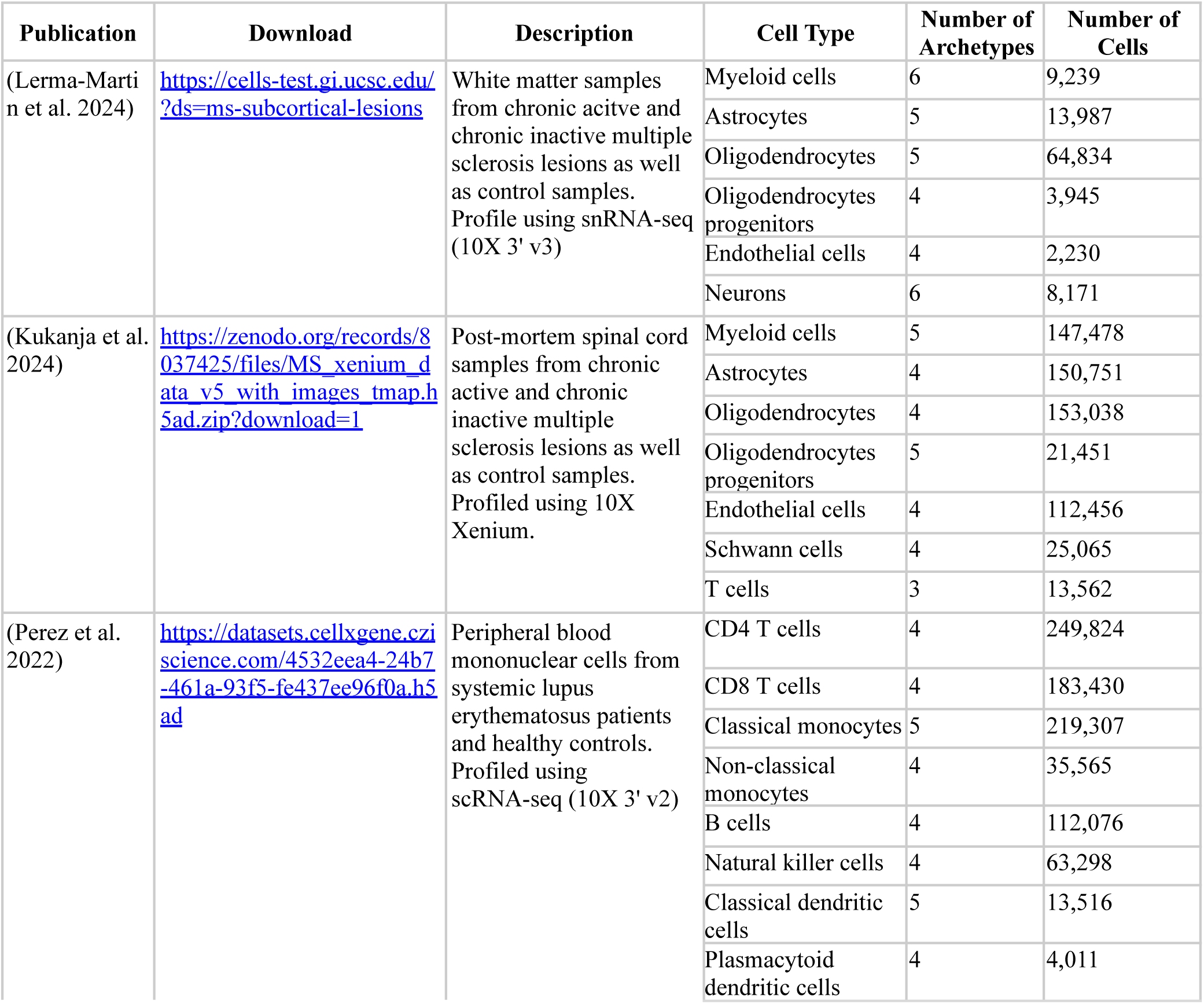
Datasets used to benchmark optimization and initialization algorithms, as well as testing different coreset sizes. Datasets: (Lerma-Martin et al. 2024) (39), (Kukanja et al. 2024) (38), (Perez et al. 2022) (40).

**Supplementary Table S2.** Datasets used to compare ParTIpy to the ParTI MATLAB package. Datasets: (Ben-Moshe et al. 2022) (29), (Chaffin et al. 2022) (37), (Kukanja et al. 2024) (38).

| Publication | Download | Description | Number of Archetypes | Number of Cells |
| --- | --- | --- | --- | --- |
| (Ben-Moshe et al. 2022) | https://zenodo.org/records/6035873<br>Files: Single_cell_Meta_data.txt, Single_cell_UMI_COUNT.txt | Murine hepatocytes profiled using scRNA-seq (10X 3' v3). We used only the hepatocytes from time point 0h. | 4 | 1,999 |
| (Chaffin et al. 2022) | https://singlecell.broadinstitute.org/single_cell/study/SCP1303/single-nuclei-profiling-of-human-dilated-and-hypertrophic-cardiomyopathy | Fibroblasts from snRNA-seq (10X 3' v3) of heart tissue from hypertrophic cardiomyopathy (HCM) patients. | 5 | 57,644 |
|  | File:<br>human_dcm_hcm_scportal_03.17.2022.h5ad |  |  |  |
| (Kukanja et al. 2024) | https://zenodo.org/records/8037425/<br>File:<br>MS_xenium_data_v5_with_images_tmap.h5ad.zip | Myeloid cells from post-mortem spinal cord samples from chronic Active and chronic inactive multiple sclerosis lesions as well as control samples profiled using 10X Xenium. | 5 | 147,478 |

## Footnotes

1 There are a few other assumptions beyond Pareto optimality that are explained in (Alon 2020; Shoval et al. 2012).

2 Shape built from straight lines and flat surfaces, like triangles and tetrahedra, extended into any number of dimensions.

## References

1. U. Alon, An Introduction to Systems Biology: Design Principles of Biological Circuits (CRC Press, Taylor & Francis Group, Boca Raton London New York, Second edition., 2020).

2. O. Shoval, H. Sheftel, G. Shinar, Y. Hart, O. Ramote, A. Mayo, E. Dekel, K. Kavanagh, U. Alon, Evolutionary Trade-Offs, Pareto Optimality, and the Geometry of Phenotype Space. Science 336, 1157–1160 (2012).

3. Y. Hart, H. Sheftel, J. Hausser, P. Szekely, N. B. Ben-Moshe, Y. Korem, A. Tendler, A. E. Mayo, U. Alon, Inferring biological tasks using Pareto analysis of high-dimensional data. Nat. Methods 12, 233–235 (2015).

4. Y. Korem, P. Szekely, Y. Hart, H. Sheftel, J. Hausser, A. Mayo, M. E. Rothenberg, T. Kalisky, U. Alon, Geometry of the Gene Expression Space of Individual Cells. PLOS Comput. Biol. 11, e1004224 (2015).

5. M. Adler, Y. Korem Kohanim, A. Tendler, A. Mayo, U. Alon, Continuum of Gene-Expression Profiles Provides Spatial Division of Labor within a Differentiated Cell Type. Cell Syst. 8, 43–52.e5 (2019).

6. M. Adler, N. Moriel, A. Goeva, I. Avraham-Davidi, S. Mages, T. S. Adams, N. Kaminski, E. Z. Macosko, A. Regev, R. Medzhitov, M. Nitzan, Emergence of division of labor in tissues through cell interactions and spatial cues. Cell Rep. 42, 112412 (2023).

7. G. Crowley, U. Alon, S. R. Quake, An Atlas of Cellular Archetypes. Molecular Biology [Preprint] (2024). 10.1101/2024.12.04.626890.

8. A. Cutler, L. Breiman, Archetypal analysis. Technometrics J. Stat. Phys. Chem. Eng. Sci. 36, 338–347 (1994).

9. M. Mørup, L. K. Hansen, Archetypal analysis for machine learning and data mining. Neurocomputing 80, 54–63 (2012).

10. A. Alcacer, I. Epifanio, S. Mair, M. Mørup, A Survey on Archetypal Analysis. arXiv arXiv:2504.12392 [Preprint] (2025). 10.48550/arXiv.2504.12392.

11. J. C. Thøgersen, M. Mørup, S. Damkiær, S. Molin, L. Jelsbak, Archetypal analysis of diverse Pseudomonas aeruginosatranscriptomes reveals adaptation in cystic fibrosis airways. BMC Bioinformatics 14, 279 (2013).

12. S. Miyara, M. Adler, K. B. Umansky, D. Häußler, E. Bassat, Y. Divinsky, J. Elkahal, D. Kain, D. Lendengolts, R. O. R. Flores, H. Bueno-Levy, O. Golani, T. Shalit, M. Gershovits, E. Weizman, A. Genzelinakh, D. M. Kimchi, A. Shakked, L. Zhang, J. Wang, A. Baehr, Z. Petrover, R. Sarig, T. Dorn, A. Moretti, J. Saez-Rodriguez, C. Kupatt, E. M. Tanaka, R. Medzhitov, A. Krüger, A. Mayo, U. Alon, E. Tzahor, Cold and hot fibrosis define clinically distinct cardiac pathologies. Cell Syst. 16 (2025).

13. S. Mohammadi, V. Ravindra, D. F. Gleich, A. Grama, A geometric approach to characterize the functional identity of single cells. Nat. Commun. 9, 1516 (2018).

14. S. M. Groves, G. V. Ildefonso, C. O. McAtee, P. M. M. Ozawa, A. S. Ireland, P. E. Stauffer, P. T. Wasdin, X. Huang, Y. Qiao, J. S. Lim, J. Bader, Q. Liu, A. J. Simmons, K. S. Lau, W. T. Iams, D. P. Hardin, E. B. Saff, W. R. Holmes, D. R. Tyson, C. M. Lovly, J. C. Rathmell, G. Marth, J. Sage, T. G. Oliver, A. M. Weaver, V. Quaranta, Archetype tasks link intratumoral heterogeneity to plasticity and cancer hallmarks in small cell lung cancer. Cell Syst. 13, 690–710.e17 (2022).

15. D. P. Cook, J. L. Wrana, A specialist-generalist framework for epithelial-mesenchymal plasticity in cancer. Trends Cancer 8, 358–368 (2022).

16. J. Hausser, U. Alon, Tumour heterogeneity and the evolutionary trade-offs of cancer. Nat. Rev. Cancer 20, 247–257 (2020).

17. A. El Marrahi, F. Lipreri, Z. Kang, L. Gsell, A. Eroglu, D. Alber, J. Hausser, NIPMAP: niche-phenotype mapping of multiplex histology data by community ecology. Nat. Commun. 14, 7182 (2023).

18. D. Túrós, J. Vasiljevic, K. Hahn, S. Rottenberg, A. Valdeolivas, Chrysalis: decoding tissue compartments in spatial transcriptomics with archetypal analysis. *Commun*. Biol. 7, 1520 (2024).

19. V. Kleshchevnikov, A. Shmatko, E. Dann, A. Aivazidis, H. W. King, T. Li, R. Elmentaite, A. Lomakin, V. Kedlian, A. Gayoso, M. S. Jain, J. S. Park, L. Ramona, E. Tuck, A. Arutyunyan, R. Vento-Tormo, M. Gerstung, L. James, O. Stegle, O. A. Bayraktar, Cell2location maps fine-grained cell types in spatial transcriptomics. Nat. Biotechnol. 40, 661–671 (2022).

20. S. He, Y. Jin, A. Nazaret, L. Shi, X. Chen, S. Rampersaud, B. S. Dhillon, I. Valdez, L. E. Friend, J. L. Fan, C. Y. Park, R. L. Mintz, Y.-H. Lao, D. Carrera, K. W. Fang, K. Mehdi, M. Rohde, J. L. McFaline-Figueroa, D. Blei, K. W. Leong, A. Y. Rudensky, G. Plitas, E. Azizi, Starfysh integrates spatial transcriptomic and histologic data to reveal heterogeneous tumor–immune hubs. Nat. Biotechnol., 1–13 (2024).

21. C. Bauckhage, K. Kersting, F. Hoppe, C. Thurau, “Archetypal analysis as an autoencoder” (2015; https://www.ml.informatik.tu-darmstadt.de/papers/autoencode2015nc2.pdf).

22. S. Mair, U. Brefeld, “Coresets for Archetypal Analysis” in Advances in Neural Information Processing Systems (Curran Associates, Inc., 2019; https://papers.nips.cc/paper_files/paper/2019/hash/7f278ad602c7f47aa76d1bfc90f20263-Abstract.html)vol. 32.

23. S. Mair, J. Sjölund, Archetypal Analysis++: Rethinking the Initialization Strategy. arXiv arXiv:2301.13748 [Preprint] (2024). 10.48550/arXiv.2301.13748.

24. I. Virshup, D. Bredikhin, L. Heumos, G. Palla, G. Sturm, A. Gayoso, I. Kats, M. Koutrouli, B. Berger, D. Pe’er, A. Regev, S. A. Teichmann, F. Finotello, F. A. Wolf, N. Yosef, O. Stegle, F. J. Theis, The scverse project provides a computational ecosystem for single-cell omics data analysis. Nat. Biotechnol. 41, 604–606 (2023).

25. A. Suleman, “Validation of archetypal analysis” in 2017 IEEE International Conference on Fuzzy Systems (FUZZ-IEEE) (2017; https://ieeexplore.ieee.org/document/8015385), pp. 1–6.

26. P. Badia-i-Mompel, J. Vélez Santiago, J. Braunger, C. Geiss, D. Dimitrov, S. Müller-Dott, P. Taus, A. Dugourd, C. H. Holland, R. O. Ramirez Flores, J. Saez-Rodriguez, decoupleR: ensemble of computational methods to infer biological activities from omics data. Bioinforma. Adv. 2, vbac016 (2022).

27. D. Dimitrov, P. S. L. Schäfer, E. Farr, P. Rodriguez-Mier, S. Lobentanzer, P. Badia-i-Mompel, A Dugourd, J. Tanevski, R. O. Ramirez Flores, J. Saez-Rodriguez, LIANA+ provides an all-in-one framework for cell–cell communication inference. Nat. Cell Biol. 26, 1613–1622 (2024).

28. D. Dimitrov, D. Türei, M. Garrido-Rodriguez, P. L. Burmedi, J. S. Nagai, C. Boys, R. O. Ramirez Flores, H. Kim, B. Szalai, I. G. Costa, A. Valdeolivas, A. Dugourd, J. Saez-Rodriguez, Comparison of methods and resources for cell-cell communication inference from single-cell RNA-Seq data. Nat. Commun. 13, 3224 (2022).

29. S. Ben-Moshe, T. Veg, R. Manco, S. Dan, D. Papinutti, A. Lifshitz, A. A. Kolodziejczyk, K. Bahar Halpern, E. Elinav, S. Itzkovitz, The spatiotemporal program of zonal liver regeneration following acute injury. Cell Stem Cell 29, 973–989.e10 (2022).

30. K. B. Halpern, R. Shenhav, O. Matcovitch-Natan, B. Tóth, D. Lemze, M. Golan, E. E. Massasa, S. Baydatch, S. Landen, A. E. Moor, A. Brandis, A. Giladi, A. Stokar-Avihail, E. David, I. Amit, S. Itzkovitz, Single-cell spatial reconstruction reveals global division of labour in the mammalian liver. Nature 542, 352–356 (2017).

31. M. Gillespie, B. Jassal, R. Stephan, M. Milacic, K. Rothfels, A. Senff-Ribeiro, J. Griss, C. Sevilla, L. Matthews, C. Gong, C. Deng, T. Varusai, E. Ragueneau, Y. Haider, B. May, V. Shamovsky, J. Weiser, T. Brunson, N. Sanati, L. Beckman, X. Shao, A. Fabregat, K. Sidiropoulos, J. Murillo, G. Viteri, J. Cook, S. Shorser, G. Bader, E. Demir, C. Sander, R. Haw, G. Wu, L. Stein, H. Hermjakob, P. D’Eustachio, The reactome pathway knowledgebase 2022. Nucleic Acids Res. 50, D687–D692 (2022).

32. J. Zhang, A. A. Ubas, R. de Borja, V. Svensson, N. Thomas, N. Thakar, I. Lai, A. Winters, U. Khan, M. G. Jones, V. Tran, J. Pangallo, E. Papalexi, A. Sapre, H. Nguyen, O. Sanderson, M. Nigos, O. Kaplan, S. Schroeder, B. Hariadi, S. Marrujo, C. C. A. Salvino, G. G. Olivares, R. Koehler, G. Geiss, A. Rosenberg, C. Roco, D. Merico, N. Alidoust, H. Goodarzi, J. Yu, Tahoe-100M: A Giga-Scale Single-Cell Perturbation Atlas for Context-Dependent Gene Function and Cellular Modeling. bioRxiv [Preprint] (2025). 10.1101/2025.02.20.639398.

33. S. M. Keller, M. Samarin, M. Wieser, V. Roth, Deep Archetypal Analysis. arXiv arXiv:1901.10799 [Preprint] (2020). 10.48550/arXiv.1901.10799.

34. S. Milite, G. Caravagna, A. Sottoriva, MIDAA: deep archetypal analysis for interpretable multi-omic data integration based on biological principles. Genome Biol. 26, 90 (2025).

35. D. van Dijk, D. B. Burkhardt, M. Amodio, A. Tong, G. Wolf, S. Krishnaswamy, “Finding Archetypal Spaces Using Neural Networks” in 2019 *IEEE International Conference on Big Data (Big Data)* (2019; https://ieeexplore.ieee.org/document/9006484), pp. 2634–2643.

36. Y. Wang, H. Zhao, Non-linear archetypal analysis of single-cell RNA-seq data by deep autoencoders. PLOS Comput. Biol. 18, e1010025 (2022).

37. M. Chaffin, I. Papangeli, B. Simonson, A.-D. Akkad, M. C. Hill, A. Arduini, S. J. Fleming, M. Melanson, S. Hayat, M. Kost-Alimova, O. Atwa, J. Ye, K. C. Bedi, M. Nahrendorf, V. K. Kaushik, C. M. Stegmann, K. B. Margulies, N. R. Tucker, P. T. Ellinor, Single-nucleus profiling of human dilated and hypertrophic cardiomyopathy. Nature 608, 174–180 (2022).

38. P. Kukanja, C. M. Langseth, L. A. Rubio Rodríguez-Kirby, E. Agirre, C. Zheng, A. Raman, C. Yokota, C. Avenel, K. Tiklová, A. O. Guerreiro-Cacais, T. Olsson, M. M. Hilscher, M. Nilsson, G. Castelo-Branco, Cellular architecture of evolving neuroinflammatory lesions and multiple sclerosis pathology. Cell 187, 1990–2009.e19 (2024).

39. C. Lerma-Martin, P. Badia-i-Mompel, R. O. R. Flores, P. Sekol, P. S. L. Schäfer, C. J. Riedl, A. Hofmann, T. Thäwel, F. Wünnemann, M. A. Ibarra-Arellano, T. Trobisch, P. Eisele, D. Schapiro, M. Haeussler, S. Hametner, J. Saez-Rodriguez, L. Schirmer, Cell type mapping reveals tissue niches and interactions in subcortical multiple sclerosis lesions. Nat. Neurosci., 1–12 (2024).

40. R. K. Perez, M. G. Gordon, M. Subramaniam, M. C. Kim, G. C. Hartoularos, S. Targ, Y. Sun, A. Ogorodnikov, R. Bueno, A. Lu, M. Thompson, N. Rappoport, A. Dahl, C. M. Lanata, M. Matloubian, L. Maliskova, S. S. Kwek, T. Li, M. Slyper, J. Waldman, D. Dionne, O. Rozenblatt-Rosen, L. Fong, M. Dall’Era, B. Balliu, A. Regev, J. Yazdany, L. A. Criswell, N. Zaitlen, C. J. Ye, Single-cell RNA-seq reveals cell type–specific molecular and genetic associations to lupus. Science 376, eabf1970 (2022).

