## Supplementary Methods for "ParTIpy: A Scalable Framework for Archetypal Analysis and Pareto Task Inference"

August 2025

### 1 Notation

- $N \in \mathbb{N}$ — number of samples (here: cells)
- $D \in \mathbb{N}$ — number of dimensions
- $G \in \mathbb{N}$ — number of genes
- $K \in \mathbb{N}$ with $K \leq N$ — number of archetypes
- $\mathcal{X} = \{\mathbf{x}_1, \ldots, \mathbf{x}_N\}$ with $\mathbf{x}_n \in \mathbb{R}^D$ — embedded coordinates of all cells
- $\mathbf{X} \in \mathbb{R}^{N \times D}$ — data matrix whose $n$'th row is $\mathbf{x}_n^T$
- $\mathcal{Z} = \{\mathbf{z}_1, \ldots, \mathbf{z}_K\}$ with $\mathbf{z}_k \in \mathbb{R}^D$ — coordinates of the $K$ archetypes
- $\mathbf{Z} \in \mathbb{R}^{K \times D}$ — archetype matrix whose $k$'th row is $\mathbf{z}_k^T$
- $\mathbf{A} \in \mathbb{R}^{N \times K}$ — coefficients that define each sample as convex combinaton of archetypes
- $\mathbf{B} \in \mathbb{R}^{K \times N}$ — coefficients that define each archetype as convex combination of samples
- $\mathbf{Y}^{(\mathbf{X})} \in \mathbb{R}^{N \times G}$ — observed gene expression matrix; row $n$ corresponds to cell $n$
- $\mathbf{Y}^{(\mathbf{Z})} \in \mathbb{R}^{K \times G}$ — inferred archetype gene expression matrix; row $k$ corresponds to archetype $k$
- $\mathbf{1}_K \in \mathbb{R}^K$ — column vector of ones
- $\mathbf{I}_K \in \mathbb{R}^{K \times K}$ — $K \times K$ identity matrix
- $\mathbb{1}[]$ — indicator function

### 2 Archetypal Analysis

#### 2.1 Objective

Let $\mathcal{X} = \{\mathbf{x}_1, \ldots, \mathbf{x}_N\}_{n=1}^N$ be a data set comprising $N$ $D$-dimensional data points, and let $\mathbf{X} \in \mathbb{R}^{N\times D}$ be the matrix where each row is a data point.

In Archetypal Analysis we make two assumptions:

1. Each data point is a convex combination of $K$ archetypes;
2. Each archetype is a convex combination of $N$ data points.

Expressing the first assumption in matrix notation yields

$$\hat{\mathbf{X}} = \mathbf{AZ} \quad \text{or} \quad \hat{\mathbf{x}}_n = \mathbf{Z}^T\mathbf{a_n} \text{ for } n = 1, ..., N \tag{1}$$

where $\hat{\mathbf{X}} \in \mathbb{R}^{N\times D}$ is the reconstructed data matrix, $\mathbf{Z} \in \mathbb{R}^{K\times D}$ is the matrix of archetypes (i.e. each row is one archetype), and $\mathbf{A} \in \mathbb{R}^{N\times K}$ is a row-stochastic matrix that defines by which archetypes each data point is formed.

Expressing the second assumption in matrix notation yields

$$\mathbf{Z} = \mathbf{BX} \quad \text{or} \quad \mathbf{z}_k = \mathbf{X}^T\mathbf{b_k} \text{ for } k = 1, ..., K \tag{2}$$

where $\mathbf{B} \in \mathbb{R}^{K\times N}$ is a row-stochastic matrix specifying the contribution of each data point to the construction of each archetype.

Here, we will quantify the reconstruction error using the RSS, given by the squared Frobenius norm,

$$\|\mathbf{X} - \hat{\mathbf{X}}\|_F^2 = \|\mathbf{X} - \mathbf{AZ}\|_F^2 = \|\mathbf{X} - \mathbf{ABX}\|_F^2 \tag{3}$$

which yields the following optimization objective

$$\begin{aligned} \hat{\mathbf{A}}, \hat{\mathbf{B}} = \underset{\substack{\mathbf{A}\in\mathbb{R}^{N\times K} \\ \mathbf{B}\in\mathbb{R}^{K\times N}}}{\arg\min} \|\mathbf{X} - \mathbf{ABX}\|_F^2 \quad \text{subject to} \\ \mathbf{A} \geq 0, \quad \mathbf{A1}_K = \mathbf{1}_N \\ \mathbf{B} \geq 0, \quad \mathbf{B1}_N = \mathbf{1}_K \end{aligned} \tag{4}$$

Introducing the set of row-stochastic matrices,

$$F(N, K) := \left\{\mathbf{A} \in \mathbb{R}^{N\times K} \mid \mathbf{A} \geq 0 \wedge \mathbf{A1}_K = \mathbf{1}_N\right\} \tag{5}$$

we can write the objective compactly as:

$$\hat{\mathbf{A}}, \hat{\mathbf{B}} = \underset{\substack{\mathbf{A}\in F(N,K) \\ \mathbf{B}\in F(K,N)}}{\arg\min} \|\mathbf{X} - \mathbf{ABX}\|_F^2 \tag{6}$$

#### 2.2 Properties of the Objective

##### 2.2.1 Translation Invariance

The minimizers $\mathbf{A}, \mathbf{B}$ of the objective are invariant under row-wise translations of $\mathbf{X}$.

Let $\tilde{\mathbf{X}} = \mathbf{X} + \mathbf{1}_N \mathbf{v}^T$ for any $\mathbf{v} \in \mathbb{R}^D$, then

$$\underset{\substack{\mathbf{A}\in F(N,K)\\ \mathbf{B}\in F(K,N)}}{\arg\min} \|\tilde{\mathbf{X}} - \mathbf{A}\mathbf{B}\tilde{\mathbf{X}}\|_F^2 = \underset{\substack{\mathbf{A}\in F(N,K)\\ \mathbf{B}\in F(K,N)}}{\arg\min} \|\mathbf{X} - \mathbf{A}\mathbf{B}\mathbf{X}\|_F^2 \tag{7}$$

For completeness, we provide our own derivation of this result in Section 8.1, following the approach of Mørup and Hansen (2012).

##### 2.2.2 Scale Invariance

The minimizers $\mathbf{A}, \mathbf{B}$ of the objective are invariant under global scaling of $\mathbf{X}$. Let $\tilde{\mathbf{X}} = \lambda \mathbf{X}$ for any $\lambda \neq 0$, then

$$\underset{\substack{\mathbf{A}\in F(N,K)\\ \mathbf{B}\in F(K,N)}}{\arg\min} \|\tilde{\mathbf{X}} - \mathbf{A}\mathbf{B}\tilde{\mathbf{X}}\|_F^2 = \underset{\substack{\mathbf{A}\in F(N,K)\\ \mathbf{B}\in F(K,N)}}{\arg\min} \|\mathbf{X} - \mathbf{A}\mathbf{B}\mathbf{X}\|_F^2 \tag{8}$$

For completeness, we provide our own derivation of this result in Section 8.2, following the approach of Mørup and Hansen (2012).

##### 2.2.3 Uniqueness up to Permutation of Archetypes

Assuming that for each archetype $k$, there exists one data point $n$ that is best reconstructed using only this archetype, i.e.

$$\forall k \in \{1, ..., K\} \exists n \in \{1, ..., N\} \; a_{nk} = 1 \tag{9}$$

and that for each archetype there exists one data point that is only used to define this archetype and not any other archetype, i.e.

$$\forall k \in \{1, ..., K\} \exists n \in \{1, ..., N\} \; b_{kn} > 0 \; \wedge \; b_{k'n} = 0 \forall k' \neq k \tag{10}$$

then the archetypes are identifiable up to permutation (i.e. the objective does not suffer from rotational ambiguity).

Formally, let $\mathbf{A} \in F(N, K)$ and $\mathbf{B} \in F(K, N)$ be feasible, and let the assumption in Equation 9 and Equation 10 hold. Consider any invertible $\mathbf{Q} \in \mathbb{R}^{K \times K}$ such that the transformed pair

$$\mathbf{A}' = \mathbf{A}\mathbf{Q}, \qquad \mathbf{B}' = \mathbf{Q}^\top \mathbf{B} \tag{11}$$

is also feasible, i.e. $\mathbf{A}' \in F(N, K)$ and $\mathbf{B}' \in F(K, N)$. Then $\mathbf{Q}$ must be a permutation matrix.

For completeness, we provide our own derivation of this result in Section 8.3, following the approach of Mørup and Hansen (2012).

##### 2.2.4 Convexity of Subproblems

If we measure the reconstruction error with the RSS, the objective in Eq. (4) is biconvex: it is convex in $\mathbf{A}$ when $\mathbf{B}$ is fixed and vice-versa. We prove convexity in $\mathbf{A}$; the argument for $\mathbf{B}$ is analogous. This biconvexity property is crucial for alternating optimization approaches, as it guarantees that each subproblem has a unique global optimum.

For completeness, we provide our own derivation of this result in Section 8.4, following the approach of Mørup and Hansen (2012).

##### 2.2.5 Only Samples Outside of the Archetypal Convex Hull Contribute to the Loss

The objective in Equation 4 can be expressed as a sum over individual data points

$$\|\mathbf{X} - \mathbf{ABX}\|_F^2 = \sum_{n=1}^{N} \|\mathbf{x}_n - \mathbf{Z}^T \mathbf{a}_n\|_2^2. \tag{12}$$

For fixed archetypes $\mathbf{Z}$, the optimal coefficient matrix $\mathbf{A}$ assign each data point to its Euclidean projection onto the convex hull of the archetypes $\mathbf{z}_1, ..., \mathbf{z}_K$, that is

$$\|\mathbf{X} - \mathbf{ABX}\|_F^2 = \sum_{n=1}^{N} \min_{\mathbf{q} \in \mathrm{conv}(\mathcal{Z})} \|\mathbf{x}_n - \mathbf{q}\|_2^2 \tag{13}$$

Hence, any point $\mathbf{x}_n \in \mathrm{conv}(\mathcal{Z})$ does not contribute to the loss.

#### 2.3 Optimization

While the objective in Equation 4 is non-convex, it is biconvex, meaning that it becomes convex in $\mathbf{A}$ when $\mathbf{B}$ is fixed, and vice versa (see Section 2.2.4). A common strategy for optimizing such biconvex objectives is to initialize $\mathbf{B}$ (and thereby $\mathbf{Z}$) and then alternate between solving the convex subproblem in one variable while keeping the other fixed. This alternating minimization procedure results in Algorithm 1.

**Algorithm 1** Prototypical Algorithm

1: **Input:** Data matrix $\mathbf{X} \in \mathbb{R}^{N \times D}$, number of archetypes $K$, max iterations $T$
2: **Initialize:** Archetypes $\mathbf{Z} \in \mathbb{R}^{K \times D}$
3: **for** $t = 1$ to $T$ **do**
4: Compute optimal weights $\mathbf{A} \in \mathbb{R}^{N \times K}$:

$$\mathbf{A} \leftarrow \underset{\mathbf{A} \in F(N,K)}{\arg\min} \; \|\mathbf{X} - \mathbf{A}\mathbf{Z}\|_F^2 \tag{14}$$

5: Compute optimal weights $\mathbf{B} \in \mathbb{R}^{K \times N}$:

$$\mathbf{B} \leftarrow \underset{\mathbf{B} \in F(K,N)}{\arg\min} \; \|\mathbf{X} - \mathbf{A}\mathbf{B}\mathbf{X}\|_F^2 \tag{15}$$

6: Update archetypes:

$$\mathbf{Z} \leftarrow \mathbf{B}\mathbf{X} \tag{16}$$

7: **if** convergence criterion is met **then**
8: **break**
9: **end if**
10: **end for**
11: **Return:** $\mathbf{A}, \mathbf{B}, \mathbf{Z}$

Building on the alternating scheme in Algorithm 1, we implement three solver variants—(i) regularised non-negative least squares (RNNLS) (Cutler and Breiman 1994), (ii) principal convex-hull analysis (PCHA) (Mørup and Hansen 2012), and (iii) a projection-free Frank-Wolfe (FW) update (Bauckhage et al. 2015) - each differing in how the simplex constraints are enforced. The next three subsections detail these variants. For a broader survey of alternative optimisers, the reader is referred to Alcacer et al. (2025).

###### 2.3.1 Regularized Nonnegative Least Squares (RNNLS)

Introduced by Cutler and Breiman (1994), this was the first algorithm to solve the archetypal analysis objective in Equation (4).

The authors proposed to solve the constrained optimization problems in lines 4 and 5 of Algorithm 1 using a Nonnegative Least Squares (NNLS) solver and enforcing the convexity constraints using a penalty term with regularization parameter $\lambda$. So for each sample $\mathbf{a}_1, ..., \mathbf{a}_N$ (i.e. each row in $\mathbf{A}$) we solve

$$\begin{aligned} \mathbf{a}_n &= \underset{\mathbf{a}_n \in \mathbb{R}^K}{\arg\min} \|\mathbf{x}_n - \mathbf{Z}^T \mathbf{a}_n\|_2^2 + \lambda \left(1 - \mathbf{1}_K^T \mathbf{a}_n\right)^2 \quad \text{subject to} \quad \mathbf{a}_n \geq 0 \\ &= \underset{\mathbf{a}_n \in \mathbb{R}^K}{\arg\min} \left\| \begin{bmatrix} \mathbf{x}_n \\ \sqrt{\lambda} \end{bmatrix} - \begin{bmatrix} \mathbf{Z}^T \\ \sqrt{\lambda} \mathbf{1}_K^T \end{bmatrix} \mathbf{a}_n \right\|_2^2 \quad \text{subject to} \quad \mathbf{a}_n \geq 0 \end{aligned} \tag{17}$$

Then to optimize $\mathbf{B}$, we first compute the optimal $\mathbf{Z}$ given the current $\mathbf{A}$ using a standard least squares solver, for example the one implemented in `numpy`.

$$\mathbf{Z} = \underset{\mathbf{Z} \in \mathbb{R}^{K \times D}}{\arg\min} \|\mathbf{X} - \mathbf{A}\mathbf{Z}\|_F^2. \tag{18}$$

Then for each archetype $\mathbf{b}_1, ..., \mathbf{b}_K$ (i.e. each row in $\mathbf{B}$) we solve

$$\begin{aligned}\mathbf{b}_k &= \underset{\mathbf{b}_k \in \mathbb{R}^N}{\arg\min} \|\mathbf{z}_k - \mathbf{X}^T \mathbf{b}_k\|_2^2 + \lambda \left(1 - \mathbf{1}_N^T \mathbf{b}_k\right)^2 \quad \text{subject to} \quad \mathbf{b}_k \geq 0 \\ &= \underset{\mathbf{b}_k \in \mathbb{R}^N}{\arg\min} \left\| \begin{bmatrix} \mathbf{z}_k \\ \sqrt{\lambda} \end{bmatrix} - \begin{bmatrix} \mathbf{X}^T \\ \sqrt{\lambda} \mathbf{1}_N^T \end{bmatrix} \mathbf{b}_k \right\|_2^2 \quad \text{subject to} \quad \mathbf{b}_k \geq 0\end{aligned} \tag{19}$$

##### 2.3.2 Principal Convex Hull Analysis (PCHA)

Inspired by the projected gradient method for NMF (Lin 2007) and normalization invariance approach introduced for NMF (Eggert and Korner 2004), the principal convex hull analysis (PCHA) algorithm was introduced by (Mørup and Hansen 2012) to solve the archetypal analysis objective in Equation (4).

First, we recast the optimization problem in terms of the $\ell_1$-normalization invariant variables $\tilde{a}_n$ and $\tilde{b}_k$ (called invariant because these variables won't change if one applies $\ell_1$-normalization).

$$\tilde{a}_{nk} = \frac{\max(a_{nk}, 0)}{\sum_{k''=1}^{K} \max(a_{nk''}, 0)}, \quad \tilde{b}_{kn} = \frac{\max(b_{kn}, 0)}{\sum_{n''=1}^{N} \max(b_{kn''}, 0)} \tag{20}$$

Thus our objective becomes

$$\hat{\mathbf{A}}, \hat{\mathbf{B}} = \underset{\substack{\mathbf{A} \in \mathbb{R}^{N \times K} \\ \mathbf{B} \in \mathbb{R}^{K \times N}}}{\arg\min} \|\mathbf{X} - \tilde{\mathbf{A}}\tilde{\mathbf{B}}\mathbf{X}\|_F^2 \quad \text{with} \quad \tilde{\mathbf{A}} = P_{\Delta_{K-1}}(\mathbf{A}), \tilde{\mathbf{B}} = P_{\Delta_{N-1}}(\mathbf{B}) \tag{21}$$

where we define $P_{\Delta_M}$ as the rowwise projection onto the $M-1$ simplex, i.e. for any matrix $\mathbf{H} \in \mathbb{R}^{N \times M}$ we have

$$\left[P_{\Delta_{M-1}}(\mathbf{H})\right]_{nm} = \frac{\max(\mathbf{H}_{nm}, 0)}{\sum_{m'=1}^{M} \max(\mathbf{H}_{nm'}, 0)} \tag{22}$$

Then the gradient of the RSS with respect to $\mathbf{a}_n$ is obtained via the chain rule

$$\frac{\partial \text{RSS}}{\partial \mathbf{a}_n} = \frac{\partial \text{RSS}}{\partial \tilde{\mathbf{a}}_n} \frac{\partial \tilde{\mathbf{a}}_n}{\partial \mathbf{a}_n} \tag{23}$$

The first part (see Section 8.7) is given by

$$\begin{aligned}\frac{\partial \text{RSS}}{\partial \tilde{\mathbf{a}}_n} &= 2\left(\tilde{\mathbf{a}}_n \mathbf{Z}\mathbf{Z}^T - \mathbf{x}_n \mathbf{Z}^T\right)^T \\ &= \left(\tilde{g}_n^{(\mathbf{A})}\right)^T\end{aligned} \tag{24}$$

The second part (see Section 8.8) is given by

$$\frac{\partial \tilde{\mathbf{a}}_n}{\partial \mathbf{a}_n} = \left( \frac{\left(\sum_{k''=1}^{K} \max\left(a_{nk''}, 0\right)\right) \mathbf{I}_K - \max\left(\mathbf{a}_n, 0\right) \mathbf{1}_K^T}{\left(\sum_{k''=1}^{K} \max\left(a_{nk''}\right)\right)^2} \right) \text{diag}\left[\mathbb{1}\left[a_1 \geq 0\right] \quad \dots \quad \mathbb{1}\left[a_K \geq 0\right]\right] \tag{25}$$

Together we have

$$\begin{aligned}\frac{\partial \mathrm{RSS}}{\partial \mathbf{a}_n} &= \frac{\partial \mathrm{RSS}}{\partial \tilde{a}_n}\frac{\partial \tilde{a}_n}{\partial a_n} \\ &= \left(\tilde{g}_n^{(\mathbf{A})}\right)^T \left(\frac{\left(\sum_{k''=1}^{K} \max\left(a_{nk''}, 0\right)\right) \mathbf{I}_K - \max\left(\mathbf{a}_n, 0\right) \mathbf{1}_K^T}{\left(\sum_{k''=1}^{K} \max\left(a_{nk''}\right)\right)^2}\right) \mathrm{diag}\left[\mathbb{1}\left[a_1 \geq 0\right] \quad \dots \quad \mathbb{1}\left[a_K \geq 0\right]\right]\end{aligned} \tag{26}$$

If we assume that $\mathbf{a}_n$ has been $\ell_1$ normalized in the previous iteration (which we can easily do, since $\ell_1$ normalization does not change the objective) the derivative simplifies to

$$\begin{aligned}\frac{\partial \mathrm{RSS}}{\partial \mathbf{a}_n} &= \left(\tilde{g}_n^{(\mathbf{A})}\right)^T \left(\mathbf{I}_K - a_n \mathbf{1}_K^T\right) \\ &= \left(\tilde{g}_n^{(A)}\right)^T \mathbf{I}_K - \left(\tilde{g}_n^{(A)}\right)^T a_n \mathbf{1}_K^T \\ &= \left(\tilde{g}_n^{(A)}\right)^T - \left(\sum_{k''=1}^{K} \tilde{g}_{nk''}^{(A)} a_{nk''}\right) \mathbf{1}_K^T\end{aligned} \tag{27}$$

So for a single element we have

$$\frac{\partial \mathrm{RSS}}{\partial a_{nk}} = \tilde{g}_{nk}^{(A)} - \sum_{k''=1}^{K} \tilde{g}_{nk''}^{(A)} a_{nk''}. \tag{28}$$

which is the same as in Section 2.2 of (Mørup and Hansen 2012).

The algorithm to update $\mathbf{A}$ is given in Algorithm 2. The gradient descent step sizes are determined using line search.

**Algorithm 2** Update $\mathbf{A}$ via Principal Convex Hull Analysis (PCHA)

**Input:** $\mathbf{X}$, $\tilde{\mathbf{A}}$, $\mathbf{Z}$
**Output:** Updated $\mathbf{A}$
$\text{RSS} \leftarrow \|\mathbf{X} - \tilde{\mathbf{A}}\mathbf{Z}\|_F^2$
$\mathbf{A} \leftarrow \tilde{\mathbf{A}}$
$\mu \leftarrow 1$
**for** $t = 1$ to $T$ **do**
  $\tilde{\mathbf{G}}^{(\mathbf{A})} \leftarrow 2\left(\tilde{\mathbf{A}}\mathbf{Z}\mathbf{Z}^T - \mathbf{X}\mathbf{Z}^T\right)$
  $\mathbf{G}^{(\mathbf{A})} \leftarrow \tilde{\mathbf{G}}^{(\mathbf{A})} - \left(\tilde{\mathbf{G}}^{(\mathbf{A})} \odot \tilde{\mathbf{A}}\right)\mathbf{1}_K\mathbf{1}_K^T$
  **for** $t^{(\text{line})} = 1$ to $T^{(\text{line})}$ **do** ▷ line search
    $\mathbf{A}_{\text{cand}} \leftarrow \mathbf{A} - \mu\mathbf{G}^{(\mathbf{A})}$
    $\tilde{\mathbf{A}}_{\text{cand}} \leftarrow P_{\Delta_{K-1}}(\mathbf{A}_{\text{cand}})$
    $\text{RSS}^{(\text{t})} \leftarrow \|\mathbf{X} - \tilde{\mathbf{A}}_{\text{cand}}\mathbf{Z}\|_F^2$
    **if** $\text{RSS}^{(\text{t})} \leq (1+\epsilon)\cdot\text{RSS}$ **then**
      $\mathbf{A} \leftarrow \mathbf{A}_{\text{cand}}$
      $\mu \leftarrow 1.2 \cdot \mu$
      $\text{RSS} \leftarrow \text{RSS}^{(\text{t})}$
      **break**
    **else**
      $\mu \leftarrow 0.5 \cdot \mu$
    **end if**
  **end for**
**end for**
**Return:** $\tilde{\mathbf{A}}$

Similarly, under the assumption that $\mathbf{b}_k$ has been $\ell_1$ normalized in the previous iteration, the gradient with respect to $\mathbf{b}_k$ is given by:

$$\frac{\partial \text{RSS}}{\partial b_k} = \left(\tilde{g}_k^{(B)}\right)^T \mathbf{I}_N - \left(\sum_{n''=1}^{N} \tilde{g}_{kn''}^{(B)} b_{kn''}\right)\mathbf{1}_N^T \tag{29}$$

The algorithm to update $\mathbf{B}$ is given in Algorithm 3.

**Algorithm 3** Update $\mathbf{B}$ via Principal Convex Hull Analysis (PCHA)

**Input:** $\mathbf{X}$, $\tilde{\mathbf{A}}$, $\tilde{\mathbf{B}}$
**Output:** Updated $\mathbf{B}$
$\mathrm{RSS} \leftarrow \|\mathbf{X} - \tilde{\mathbf{A}}\tilde{\mathbf{B}}\mathbf{X}\|_F^2$
$\mathbf{B} \leftarrow \tilde{\mathbf{B}}$
$\mu \leftarrow 1$
**for** $t = 1$ to $T$ **do**
  $\tilde{\mathbf{G}}^{(\mathbf{B})} \leftarrow 2\left(\tilde{\mathbf{A}}^T\tilde{\mathbf{A}}\tilde{\mathbf{B}}\mathbf{X}\mathbf{X}^T - \tilde{\mathbf{A}}^T\mathbf{X}\mathbf{X}^T\right)$
  $\mathbf{G}^{(\mathbf{B})} \leftarrow \tilde{\mathbf{G}}^{(\mathbf{B})} - \left(\tilde{\mathbf{G}}^{(\mathbf{B})} \odot \tilde{\mathbf{B}}\right)\mathbf{1}_N\mathbf{1}_N^T$
  **for** $t^{(\text{line})} = 1$ to $T^{(\text{line})}$ **do** ▷ line search
    $\mathbf{B}_{\text{cand}} \leftarrow \mathbf{B} - \mu\mathbf{G}^{(\mathbf{B})}$
    $\tilde{\mathbf{B}}_{\text{cand}} \leftarrow P_{\Delta_{N-1}}(\mathbf{B}_{\text{cand}})$
    $\mathrm{RSS}^{(t)} \leftarrow \|\mathbf{X} - \tilde{\mathbf{A}}\tilde{\mathbf{B}}_{\text{cand}}\mathbf{X}\|_F^2$
    **if** $\mathrm{RSS}^{(t)} \leq (1+\epsilon)\cdot\mathrm{RSS}$ **then**
      $\mathbf{B} \leftarrow \tilde{\mathbf{B}}_{\text{cand}}$
      $\mu \leftarrow 1.2\cdot\mu$
      $\mathrm{RSS} \leftarrow \mathrm{RSS}^{(t)}$
      **break**
    **else**
      $\mu \leftarrow 0.5\cdot\mu$
    **end if**
  **end for**
**end for**
**Return:** $\tilde{\mathbf{B}}$

##### 2.3.3 Frank-Wolfe (FW) Algorithm

The Frank-Wolfe (FW) algorithm is a first-order iterative method for solving constrained optimization problems. In contrast to projected-gradient methods, such as PCHA, FW replaces the projection step with a linear minimization over the feasible set. Specifically, it solves a linear approximation of the objective at each iterate and moves toward the minimizer, thereby ensuring that all updates remain within the feasible region. For the classical FW algorithm to be applicable, the feasible set must be compact and convex. The absence of projections makes FW particularly attractive for structured domains where linear minimization is computationally cheaper than projection. For more details, see (Bauckhage et al. 2015; Clarkson 2010; Frank and Wolfe 1956; Jaggi 2013).

In our alternating optimization scheme the optimization in $\mathbf{A}$ (resp. $\mathbf{B}$) is over the standard $(K-1)$-simplex (resp. $(N-1)$-simplex), both of which are convex (Appendix 8.5) and compact (Appendix 8.6). Thus the prerequisites for the vanilla FW algorithm are met.

Given a continuously differentiable objective function $f : \mathcal{D} \to \mathbb{R}$, and convex, compact feasible set $\mathcal{D}$, the FW algorithm is outlined in Algorithm 4.

**Algorithm 4** Vanilla Frank-Wolfe (Frank and Wolfe 1956; Jaggi 2013)

1: **Input:** Convex, compact feasible set $\mathcal{D}$, continuously differentiable objective function $f : \mathcal{D} \to \mathbb{R}$, number of iterations $T$
2: **Initialize:** $\mathbf{x}^{(1)} \in \mathcal{D}$
3: **for** $t = 1$ to $T$ **do**
4: Compute linear minimizer:
$$\mathbf{s}^{(t)} = \underset{\mathbf{s} \in \mathcal{D}}{\arg\min} \left(\nabla f\left(\mathbf{x}^{(t)}\right)\right)^T \mathbf{s} \tag{30}$$
5: Set step size:
$$\mu^{(t)} = \frac{2}{t+2} \tag{31}$$
6: Update $\mathbf{x}$:
$$\mathbf{x}^{(t+1)} = (1 - \mu^{(t)})\mathbf{x}^{(t)} + \mu^{(t)}\mathbf{s}^{(t)} \tag{32}$$
7: **end for**
8: **Return:** $\mathbf{x}^{(T+1)}$

Because $0 \leq \mu^{(t)} \leq 1$ and both $\mathbf{x}^{(t)}$ and $\mathbf{s}^{(t)}$ lie in $\mathcal{D}$, every iterate $x^{(t+1)}$ remains in the feasible set by convexity.

Now, in the case of archetypal analysis, when updating the rows of $\mathbf{A}$, the feasible set is the standard simplex $\Delta_{K-1}$:
$$\mathbf{a}_n = \underset{\mathbf{a}_n \in \Delta_{K-1}}{\arg\min} \|\mathbf{x}_n - \mathbf{Z}^T \mathbf{a}_n\|_2^2. \tag{33}$$

As shown in Equation 122, the gradient with respect to $\mathbf{a}_n$ is given by
$$g_n^{(\mathbf{A})} = 2\left(\mathbf{a}_n^T \mathbf{Z}\mathbf{Z}^T - \mathbf{x}_n^T \mathbf{Z}\right). \tag{34}$$

Accordingly, the FW linear subproblem becomes
$$\mathbf{s}^{(t)} = \underset{\mathbf{s} \in \Delta_{K-1}}{\arg\min} \left(g_n^{(\mathbf{A})}\right)^T \mathbf{s}. \tag{35}$$

Since the objective is linear in $\mathbf{s}$ and $\Delta_{K-1}$ is convex and compact, the minimizer is attained at a vertex of the simplex. That is, the solution lies in $\{\mathbf{e}_1, \ldots, \mathbf{e}_K\}$, where $\mathbf{e}_k$ denotes the $k$th standard basis vector:
$$\mathbf{s}^{(t)} = \underset{\mathbf{s} \in \{\mathbf{e}_1, \ldots, \mathbf{e}_K\}}{\arg\min} \left(g_n^{(\mathbf{A})}\right)^T \mathbf{s}. \tag{36}$$

This simply selects the direction of steepest descent, i.e. the component where the gradient is most negative:
$$\mathbf{s}^{(t)} = \mathbf{e}_{k'} \quad \text{where} \quad k' = \underset{k \in \{1, \ldots, K\}}{\arg\min}\, g_{nk}^{(\mathbf{A})}. \tag{37}$$

Putting everything together, we obtain the update rule for $\mathbf{A}$, as outlined in Algorithm 5.

**Algorithm 5** Update A via Frank-Wolfe

**Input:** $\mathbf{X}$, $\mathbf{A}$, $\mathbf{Z}$
**Output:** Updated $\mathbf{A}$
Initialize: $\mathbf{A}^{(1)} = \mathbf{A}$
**for** $t = 1$ to $T$ **do**
  Compute gradient for all rows of $\mathbf{A}$:

$$\mathbf{G}^{(\mathbf{A})} \leftarrow 2\left(\mathbf{A}\mathbf{Z}\mathbf{Z}^T - \mathbf{X}\mathbf{Z}^T\right) \tag{38}$$

  **for** $n = 1$ to $N$ **do**
    Compute linear minimizer:

$$\mathbf{s}_n^{(t)} = \mathbf{e}_{k'} \quad \text{where} \quad k' = \arg\min_{k \in \{1,\dots,K\}} g_{nk}^{(\mathbf{A})} \tag{39}$$

    Set step size:

$$\mu^{(t)} = \frac{2}{t+2} \tag{40}$$

    Update $\mathbf{a}_n$:

$$\mathbf{a}_n^{(t+1)} = (1 - \mu^{(t)})\mathbf{a}_n^{(t)} + \mu^{(t)}\mathbf{s}_n^{(t)} \tag{41}$$

  **end for**
**end for**
**Return:** $\mathbf{A}^{(T+1)}$

Following the derivation of the FW update of **A**, the FW update for **B** is outlined in Algorithm 6.

**Algorithm 6** Update B via Frank-Wolfe

**Input:** $\mathbf{X}$, $\mathbf{A}$, $\mathbf{B}$
**Output:** Updated $\mathbf{B}$
Initialize: $\mathbf{B}^{(1)} = \mathbf{B}$
**for** $t = 1$ to $T$ **do**
  Compute gradient for all rows of $\mathbf{B}$:

$$\mathbf{G}^{(\mathbf{B})} \leftarrow 2\left(\mathbf{A}^T\mathbf{A}\mathbf{B}\mathbf{X}\mathbf{X}^T - \mathbf{A}^T\mathbf{X}\mathbf{X}^T\right) \tag{42}$$

  **for** $k = 1$ to $K$ **do**
    Compute linear minimizer:

$$\mathbf{s}_k^{(t)} = \mathbf{e}_{n'} \quad \text{where} \quad n' = \arg\min_{n \in \{1,\dots,N\}} g_{kn}^{(\mathbf{B})} \tag{43}$$

    Set step size:

$$\mu^{(t)} = \frac{2}{t+2} \tag{44}$$

    Update $\mathbf{b}_k$:

$$\mathbf{b}_k^{(t+1)} = (1 - \mu^{(t)})\mathbf{b}_k^{(t)} + \mu^{(t)}\mathbf{s}_k^{(t)} \tag{45}$$

  **end for**
**end for**
**Return:** $\mathbf{B}^{(T+1)}$

#### 2.4 Initialization

##### 2.4.1 Uniform

Sampling data points with uniform probability from the data set to initialize the archetypes was the first initialization scheme used for archetypal analysis (Cutler and Breiman 1994). However, already this original work the authors stated that initializing archetypes too close to each other can impede convergence speed.

##### 2.4.2 Furthest Sum

Inspired by the *FurthestFirst* initialization for $K$-means clustering (Hochbaum and Shmoys 1985), the *FurthestSum* initialization for archetypal analysis was introduced by Mørup and Hansen (2012). The algorithm is outlined in Algorithm 7. The idea is to greedily select points such that each point that is added, maximizes the sum of distances to all the other points that have already been chosen. To dampen the effect of an "unlucky" random seed, we let the selection run for $K + 10$ iterations and discard the first 10 indices, this "burn-in" leaves exactly $K$ archetypes.

**Algorithm 7** Furthest Sum Initialization

**Input:** Data matrix $\mathbf{X} \in \mathbb{R}^{N \times D}$, number of archetypes $K \leq N$
**Output:** Set of selected indices $\mathcal{S}$
Define full index set $\mathcal{I} = \{1, \dots, N\}$
Initialize set of selected indices $\mathcal{S} \leftarrow (i_1)$, where $i_1$ is randomly chosen
**for** $t = 2$ to $K + 10$ **do**
    **if** $|\mathcal{S}| > K$ **then**
        $\mathcal{S} \leftarrow \mathcal{S} \setminus i_{t-K}$ ▷ Remove the oldest index from $\mathcal{S}$
    **end if**
    **for** $n \in \mathcal{I} \setminus \mathcal{S}$ **do** ▷ Compute distances to current archetypes
        $d_n^{(\text{sum})} = \sum_{s \in \mathcal{S}} \|x_n - x_s\|_2$
    **end for**
    $i_t = \arg\max_{n \in \mathcal{I} \setminus \mathcal{S}} d_n^{(\text{sum})}$ ▷ Select new data point
    $\mathcal{S} \leftarrow \mathcal{S} \cup \{i_t\}$ ▷ Add $i_t$ to $\mathcal{S}$
**end for**
**Return:** $\mathcal{S}$

##### 2.4.3 Archetypal Analysis++

The *FurthestSum* algorithm does not guarantee that the selected archetypes are non-redundant. An archetype is redundant if it lies in the convex hull of the already selected archetypes (Suleman 2017b). Furthermore, in some cases it has been reported that the *FurthestSum* initialization yields inferior results compared to the uniform initialization (Krohne et al. 2019; Olsen et al. 2022). To address these limitations, Mair and Sjölund (2024) introduced the *AA++* initialization algorithm outlined in Algorithm 8. At each iteration, the sampling probability is proportional to distance to the convex hull of the already selected archetypes, which ensures that new archetypes are not redundant.

**Algorithm 8** Archetypal++ Initialization

**Input:** Data matrix $\mathbf{X} \in \mathbb{R}^{N \times D}$, number of archetypes $K \leq N$
**Output:** Set of selected indices $\mathcal{S}$
Initialize set of selected indices $\mathcal{S} \leftarrow \{i_1\}$, where $i_1$ is randomly chosen
**for** $t = 2$ to $K$ **do**
  Let $\mathbf{Z}$ be the submatrix of $\mathbf{X}$ with rows in $\mathcal{S}$
  **for** $n = 1$ to $N$ **do**
    $d_n = \min_{a_n} \left\|x_n - \mathbf{Z}^T a_n\right\|_2$ ▷ Compute distance to convex hull of $\mathbf{Z}$
  **end for**
  **for** $n = 1$ to $N$ **do**
    $p_n = \frac{d_n}{\sum_{n'}^{N} d_{n'}}$ ▷ Compute sample probability
  **end for**
  $i_t \sim \text{Categorical}(\mathbf{p})$ ▷ Sample new data point
  $\mathcal{S} \leftarrow \mathcal{S} \cup \{i_t\}$ ▷ Add $i_t$ to $\mathcal{S}$
**end for**
**Return:** $\mathcal{S}$

#### 2.5 Coresets

The idea of a *coreset* is to replace the full dataset $\mathcal{X} = \{\mathbf{x}_n\}_{n=1}^{N}$ with a much smaller, *weighted* subset

$$\tilde{\mathcal{X}} := \{(w_{\tilde{n}} \in \mathbb{R}_+,\ \mathbf{x}_{\tilde{n}} \in \mathcal{X})\}_{\tilde{n}=1}^{\tilde{N}} \quad \text{where} \quad \tilde{N} \ll N \tag{46}$$

such that models fitted on the coreset also provide a good fit on the original dataset (Feldman 2020).

However there are many different *coreset* definitions, as reviewed by Feldman (2020). In the following, we focus on three specific classes of coresets. Consider an optimization problem of the form

$$\min_{\mathcal{Z} \in \Theta} \sum_{n=1}^{N} d(\mathbf{x}_n, \mathcal{Z})^2 \tag{47}$$

where $\Theta := \{\{z_1, ..., z_{K'}\} \subset \mathbb{R}^D \mid K' \leq K\}$ denotes the set of candidate solution sets $\mathcal{Z} \subset \mathbb{R}^D$ of cardinality at most $K$, and $d$ denotes some distance or loss function, for example in archetypal analysis (see Section 2.2.5) and $K$-means clustering:

$$d(\mathbf{x}_n, \mathcal{Z}) = \begin{cases} \min\limits_{\mathbf{q} \in \text{conv}(\mathcal{Z})} \|\mathbf{x}_n - \mathbf{q}\|_2^2, & \text{(Archetypal Analysis)} \\ \min\limits_{\mathbf{z} \in \mathcal{Z}} \|\mathbf{x}_n - \mathbf{z}\|_2^2, & \text{($K$-means)} \end{cases} \tag{48}$$

Now, for $\varepsilon \in [0, 1]$ and $K \in \mathbb{N}$, a weighted subset $\tilde{\mathcal{X}} = \{(w_{\tilde{n}}, x_{\tilde{n}})\}_{\tilde{n}=1}^{\tilde{N}}$ with $x_{\tilde{n}} \in \mathcal{X}$ and $w_{\tilde{n}} \in \mathbb{R}_+$ is called an $\varepsilon$-coreset, $\varepsilon$-lightweight-coreset, or $\varepsilon$-absolute-coreset, respectively, if for all $\mathcal{Z} \in \Theta$,

$$\left| \sum_{n=1}^{N} d(\mathbf{x}_n, \mathcal{Z}) \; - \; \sum_{\tilde{n}=1}^{\tilde{N}} w_{\tilde{n}}\, d(\mathbf{x}_{\tilde{n}}, \mathcal{Z}) \right| \;\leq\; \begin{cases} \varepsilon \sum_{n=1}^{N} d(\mathbf{x}_n, \mathcal{Z}), & \varepsilon\text{-coreset}, \\ \frac{\varepsilon}{2} \sum_{n=1}^{N} d(\mathbf{x}_n, \mathcal{Z}) + \frac{\varepsilon}{2} \sum_{n=1}^{N} d(\mathbf{x}_n, \{\bar{\mathbf{x}}\}), & \varepsilon\text{-lightweight-coreset}, \\ \varepsilon, & \varepsilon\text{-absolute-coreset}. \end{cases} \tag{49}$$

The standard (relative) $\varepsilon$-coreset is included here only as a reference, as it is the variant most widely used in the coreset literature and provides a point of comparison with the lightweight and absolute forms. It is usually written in the multiplicative form

$$\forall \mathcal{Z} \in \Theta : \quad (1-\varepsilon)\sum_{n=1}^{N} d(\mathbf{x}_n, \mathcal{Z}) \leq \sum_{\tilde{n}=1}^{\tilde{N}} w_{\tilde{n}} d(\mathbf{x}_{\tilde{n}}, \mathcal{Z}) \leq (1+\varepsilon)\sum_{n=1}^{N} d(\mathbf{x}_n, \mathcal{Z}) \tag{50}$$

which highlights the $(1 \pm \varepsilon)$ multiplicative approximation guarantees.

Mair and Brefeld (2019) showed that distance used in archetypal analysis is upper-bounded by the distance used in $K$-means clustering (see Lemma 1 and Proposition 1). Hence, the archetypal analysis loss is upper bounded by the $K$-means loss, i.e. for all $\mathcal{Z} \in \Theta$ we have

$$\sum_{n=1}^{N} \min_{\mathbf{z}\in \operatorname{conv}(\mathcal{Z})} \|\mathbf{x}_n - \mathbf{z}\|_2^2 \leq \sum_{n=1}^{N} \min_{\mathbf{z}\in \mathcal{Z}} \|\mathbf{x}_n - \mathbf{z}\|_2^2 \tag{51}$$

Consequently, any $\varepsilon$-coreset for $K$-means also satisfies Equation 49 for AA, with the same $\varepsilon$. Mair and Brefeld (2019) then constructed the first *$\varepsilon$-absolute-coreset* for archetypal analysis by adapting the *lightweight-coreset* construction of Bachem, Lucic, and Krause (2018). Note that the coreset of Mair and Brefeld (2019) is not provably valid for every $\mathcal{Z} \in \Theta$; its guarantee holds only when the data mean $\bar{\mathbf{x}}$ lies in $\operatorname{conv}(\mathcal{Z})$ (see Corollary 1). In practice this requirement is mild, because archetypes are extreme points and their convex hull typically encloses $\bar{\mathbf{x}}$. The resulting algorithm is outlined in Algorithm 9.

**Algorithm 9** Coreset Construction for Archetypal Analysis

**Input:** Data matrix $\mathbf{X} \in \mathbb{R}^{N\times D}$, coreset size $\tilde{N} < N$
**Output:** Coreset $\tilde{\mathbf{X}}$
$\bar{\mathbf{x}} \leftarrow \frac{1}{N}\sum_{n=1}^{N} \mathbf{x}_n$ ▷ Compute mean of data
**for** $n = 1$ to $N$ **do**
  $q(\mathbf{x}_n) = \frac{d(\mathbf{x}_n, \bar{\mathbf{x}})^2}{\sum_{n'=1}^{N} d(\mathbf{x}_{n'}, \bar{\mathbf{x}})^2}$ ▷ Compute sampling probability
**end for**
$\tilde{\mathcal{X}} \leftarrow \emptyset$
**for** $\tilde{n} = 1$ to $\tilde{N}$ **do**
  $\tilde{\mathbf{x}}_{\tilde{n}} \sim q(\mathbf{x}_1, ..., \mathbf{x}_N)$ ▷ Sample points with replacement
  $w_{\tilde{n}} = \frac{1}{\tilde{N} q(\tilde{\mathbf{x}}_{\tilde{n}})}$ ▷ Assign weight
  $\tilde{\mathcal{X}} \leftarrow \tilde{\mathcal{X}} \cup \{(w_{\tilde{n}}, \mathbf{x}_{\tilde{n}})\}$
**end for**
**Return:** $\tilde{\mathcal{X}} = \{(w_{\tilde{n}}, \tilde{\mathbf{x}}_{\tilde{n}})\}_{\tilde{n}=1}^{\tilde{N}}$

Accordingly the archetypal analysis objective from Equation 4 is adapted to incorporate weights

$$\begin{aligned} \mathbf{A}, \mathbf{B} &= \underset{\substack{\mathbf{A}\in F(\tilde{N},K)\\ \mathbf{B}\in F(K,\tilde{N})}}{\arg\min} \sum_{\tilde{n}=1}^{\tilde{N}} w_{\tilde{n}} \|\tilde{\mathbf{x}}_{\tilde{n}} - \tilde{\mathbf{X}}^T \mathbf{B}^T \mathbf{a}_{\tilde{n}}\|_2^2 \\ &= \underset{\substack{\mathbf{A}\in F(\tilde{N},K)\\ \mathbf{B}\in F(K,\tilde{N})}}{\arg\min} \|\mathbf{W}\tilde{\mathbf{X}} - \mathbf{WAB}\tilde{\mathbf{X}}\|_F^2 \end{aligned} \tag{52}$$

where $\mathbf{W} := \operatorname{diag}(\sqrt{w_1}, ..., \sqrt{w_{\tilde{N}}})$ and $\tilde{\mathbf{X}} \in \mathbb{R}^{\tilde{N}\times D}$ denotes the subsampled data. Because the weights do not influence the update rule for $\mathbf{A}$ (since each $\mathbf{a}_n$ is updated independently), it suffices to replace $\mathbf{X}$ by $\tilde{\mathbf{X}}$. In contrast, the update rule for $\mathbf{B}$ must be adjusted to account for the weights; it depends on both $\tilde{\mathbf{X}}$ and the weighted matrix $\mathbf{W}\tilde{\mathbf{X}}$.

$$\mathbf{B} \leftarrow \underset{\mathbf{B}\in F(K,\tilde{N})}{\arg\min} \left\|\mathbf{W}\tilde{\mathbf{X}} - \mathbf{WAB}\tilde{\mathbf{X}}\right\|_F^2 \tag{53}$$

Originally the authors adapted the regularized nonnegative least squares (see Equation 19) by recasting the update for $\mathbf{B}$ as

$$\begin{aligned}\mathbf{Z} &\leftarrow \underset{\mathbf{Z}\in\mathbb{R}^{K\times D}}{\arg\min}\|\mathbf{W}\tilde{\mathbf{X}} - \mathbf{W}\mathbf{A}\mathbf{Z}\|_F^2 \\ \mathbf{b}_k &\leftarrow \underset{\mathbf{b}_k\in\mathbb{R}^{\tilde{N}}}{\arg\min}\left\|\begin{bmatrix}\mathbf{z}_k \\ \sqrt{\lambda}\end{bmatrix} - \begin{bmatrix}\tilde{\mathbf{X}}^T \\ \sqrt{\lambda}\mathbf{1}_{\tilde{N}}^T\end{bmatrix} b_k\right\|_2^2\end{aligned} \tag{54}$$

To fully capitalize on the computational speed-ups afforded by the coreset, we replace the NNLS update with the more efficient PCHA and FW algorithm. Accordingly, we re-derive their update rules from the weighted objective in Equation 53, using its gradient with respect to $\mathbf{B}$

$$\begin{aligned}\tilde{\mathbf{G}}^{(B)} &= \nabla_B \widetilde{\mathrm{RSS}} \\ &= \nabla_B \|\mathbf{W}\tilde{\mathbf{X}} - \mathbf{W}\mathbf{A}\mathbf{B}\tilde{\mathbf{X}}\|_F^2 \\ &= 2\left((\mathbf{W}\mathbf{A})^T(\mathbf{W}\mathbf{A})\mathbf{B}\tilde{\mathbf{X}}\tilde{\mathbf{X}}^T - (\mathbf{W}\mathbf{A})^T\left(\mathbf{W}\tilde{\mathbf{X}}\right)\tilde{\mathbf{X}}^T\right)\end{aligned} \tag{55}$$

To summarize, Algorithm 10 shows how to adapt a generic archetypal analysis solver to a coreset.

**Algorithm 10** Prototypical Algorithm Optimization using Coreset

**Input:** Data matrix $\mathbf{X} \in \mathbb{R}^{N\times D}$, subsampled data matrix $\tilde{\mathbf{X}} \in \mathbb{R}^{\tilde{N}\times D}$, coreset weights $\mathbf{W} \in \mathbb{R}_+^{\tilde{N}\times\tilde{N}}$ number of archetypes $K$, max iterations $T$
**Initialize:** Archetypes $\mathbf{Z} \in \mathbb{R}^{K\times D}$
**for** $t = 1$ to $T$ **do**
  Compute optimal weights $\mathbf{A} \in \mathbb{R}^{\tilde{N}\times K}$:

$$\mathbf{A} \leftarrow \underset{\mathbf{A}\in F(\tilde{N},K)}{\arg\min}\ \|\tilde{\mathbf{X}} - \mathbf{A}\mathbf{Z}\|_F^2 \tag{56}$$

  Compute optimal weights $\mathbf{B} \in \mathbb{R}^{K\times\tilde{N}}$:

$$\mathbf{B} \leftarrow \underset{\mathbf{B}\in F(K,\tilde{N})}{\arg\min}\ \|\mathbf{W}\tilde{\mathbf{X}} - \mathbf{W}\mathbf{A}\mathbf{B}\tilde{\mathbf{X}}\|_F^2 \tag{57}$$

  Update archetypes:

$$\mathbf{Z} \leftarrow \mathbf{B}\tilde{\mathbf{X}} \tag{58}$$

  **if** convergence criterion is met **then**
    **break**
  **end if**
**end for**
Recompute $\mathbf{A} \in F(N, K)$ on full dataset

$$\mathbf{A} \leftarrow \underset{\mathbf{A}\in F(N,K)}{\arg\min}\ \|\mathbf{X} - \mathbf{A}\mathbf{Z}\|_F^2 \tag{59}$$

**Return:** $\mathbf{A}, \mathbf{B}, \mathbf{Z}$

In particular, only the following adaptions are required

1. Pre-compute the weighted data matrix, $\mathbf{W}\tilde{\mathbf{X}}$.
2. Evaluate the gradient with respect to $\mathbf{B}$ using Equation (55).
3. After the algorithm converges, optimize $\mathbf{A}$ on the full dataset $\mathbf{X}$ while holding $\mathbf{Z} = \mathbf{B}\tilde{\mathbf{X}}$ fixed.

#### 2.6 Relaxation of Archetype Constraints

We can relax the constraint that archetypes must be convex combinations of data points (i.e. that they must lie within the convex hull of the data).

$$\begin{aligned}\hat{\mathbf{A}}, \hat{\mathbf{B}} = &\operatorname*{arg\,min}_{\substack{\mathbf{A}\in\mathbb{R}^{N\times K}\\ \mathbf{B}\in\mathbb{R}^{K\times N}}} \|\mathbf{X} - \mathbf{ABX}\|_F^2 \quad \text{subject to} \\ &\mathbf{A} \geq 0, \quad \mathbf{A}\mathbf{1}_K = \mathbf{1}_N \\ &\mathbf{B} \geq 0, \quad \forall k \in \{1, ..., K\} \quad 1-\delta \leq \|\mathbf{b}_k\|_1 \leq 1+\delta\end{aligned} \tag{60}$$

Note that changing the constraint on $\mathbf{B}$ into two inequality constraints, the objective is still convex in $\mathbf{B}$ given a fixed $\mathbf{A}$ (Mørup and Hansen 2012). To optimize Equation 60, Mørup and Hansen (2012) first rewrite the objective by introducing a scaling vector $\boldsymbol{\alpha} \in \mathbb{R}_+^K$

$$\begin{aligned}\hat{\mathbf{A}}, \hat{\mathbf{B}} = &\operatorname*{arg\,min} \|\mathbf{X} - \mathbf{A}\operatorname{diag}(\boldsymbol{\alpha})\mathbf{BX}\|_F^2 \quad \text{subject to} \\ &\mathbf{A} \in F(N, K) \\ &\mathbf{B} \in F(K, N) \\ &\forall k \in \{1, ..., K\} \quad 1-\delta \leq \alpha_k \leq 1+\delta\end{aligned} \tag{61}$$

such that each row $k$ of $\mathbf{B}$ is scaled by $\alpha_k$. This formulation readily extends both the PCHA and the FW algorithm to the relaxed objective. Specifically, we can apply either method to update $\mathbf{A}$ and $\mathbf{B}$, using the gradients derived in Section 8.9 and shown below. The gradient with respect to $\mathbf{A}$ is structurally identical to the standard case, with the archetype matrix given by $\mathbf{Z} = \operatorname{diag}(\boldsymbol{\alpha})\mathbf{BX}$.

$$\begin{aligned}\mathbf{G}^{(A)} &= \nabla_{\mathbf{A}} \|\mathbf{X} - \mathbf{A}\operatorname{diag}(\boldsymbol{\alpha})\mathbf{BX}\|_F^2 \\ &= 2\left(\mathbf{A}\operatorname{diag}(\boldsymbol{\alpha})\mathbf{BXX}^T\mathbf{B}^T\operatorname{diag}(\boldsymbol{\alpha}) - \mathbf{XX}^T\mathbf{B}^T\operatorname{diag}(\boldsymbol{\alpha})\right) \\ &= 2\left(\mathbf{AZZ}^T - \mathbf{XZ}^T\right)\end{aligned} \tag{62}$$

For $\mathbf{B}$ the gradient is given by

$$\begin{aligned}\mathbf{G}^{(B)} &= \nabla_{\mathbf{B}} \|\mathbf{X} - \mathbf{A}\operatorname{diag}(\boldsymbol{\alpha})\mathbf{BX}\|_F^2 \\ &= 2\left[\operatorname{diag}(\boldsymbol{\alpha})\mathbf{A}^T\mathbf{A}\operatorname{diag}(\boldsymbol{\alpha})\mathbf{BXX}^T - \operatorname{diag}(\boldsymbol{\alpha})\mathbf{A}^T\mathbf{XX}^T\right]\end{aligned} \tag{63}$$

At each iteration, $\boldsymbol{\alpha}$ can be updated via projected gradient descent using the gradient

$$\begin{aligned}\mathbf{G}^{(\alpha)} &= \nabla_{\alpha} \|\mathbf{X} - \mathbf{A}\operatorname{diag}(\boldsymbol{\alpha})\mathbf{BX}\|_F^2 \\ &= 2\operatorname{diag}\left(\mathbf{A}^T\mathbf{A}\operatorname{diag}(\boldsymbol{\alpha})\mathbf{BXX}^T\mathbf{B}^T - \mathbf{A}^T\mathbf{XX}^T\mathbf{B}^T\right)\end{aligned} \tag{64}$$

and the component-wise projection onto $[1-\delta, 1+\delta]$.

The algorithm is outlined in Algorithm 11.

**Algorithm 11** Prototypical Algorithm for Objective with Relaxed Archetypal Constraints

**Input:** Data matrix $\mathbf{X} \in \mathbb{R}^{N \times D}$, number of archetypes $K$, max iterations $T$
**Initialize:** Archetypes $\mathbf{Z} = \mathbf{BX} \in \mathbb{R}^{K \times D}$, $\boldsymbol{\alpha} = \mathbf{1}_K$
**for** $t = 1$ to $T$ **do**
    Compute optimal weights $\mathbf{A} \in \mathbb{R}^{N \times K}$:

$$\mathbf{A} \leftarrow \underset{\mathbf{A} \in F(N,K)}{\arg\min} \; \|\mathbf{X} - \mathbf{A}\,\mathrm{diag}(\boldsymbol{\alpha})\mathbf{BX}\|_F^2 \tag{65}$$

    Compute optimal weights $\mathbf{B} \in \mathbb{R}^{K \times N}$:

$$\mathbf{B} \leftarrow \underset{\mathbf{B} \in F(K,N)}{\arg\min} \; \|\mathbf{X} - \mathbf{A}\,\mathrm{diag}(\boldsymbol{\alpha})\mathbf{BX}\|_F^2 \tag{66}$$

    Compute optimal $\boldsymbol{\alpha} \in [1-\delta, 1+\delta]^K$:

$$\boldsymbol{\alpha} \leftarrow \underset{\boldsymbol{\alpha} \in [1-\delta, 1+\delta]^K}{\arg\min} \; \|\mathbf{X} - \mathbf{A}\,\mathrm{diag}(\boldsymbol{\alpha})\mathbf{BX}\|_F^2 \tag{67}$$

    Update archetypes:

$$\mathbf{Z} \leftarrow \mathrm{diag}(\boldsymbol{\alpha})\mathbf{BX} \tag{68}$$

    **if** convergence criterion is met **then**
        **break**
    **end if**
**end for**
**Return:** $\mathbf{A}, \mathbf{B}, \mathbf{Z}$

#### 2.7 Combining Coresets and Relaxation of Archetype Constraints

If we seek to use coresets and relax the constraint that archetypes must lie within the convex hull of the data, then we need to optimize the objective

$$\begin{aligned} \hat{\mathbf{A}}, \hat{\mathbf{B}} = \arg\min & \|\mathbf{W}\tilde{\mathbf{X}} - \mathbf{WA}\,\mathrm{diag}(\boldsymbol{\alpha})\mathbf{B}\tilde{\mathbf{X}}\|_F^2 \quad \text{subject to} \\ & \mathbf{A} \in F(\tilde{N}, K) \\ & \mathbf{B} \in F(K, \tilde{N}) \\ & \forall k \in \{1, ..., K\} \quad 1 - \delta \leq \alpha_k \leq 1 + \delta \end{aligned} \tag{69}$$

As in Section 2.6, we can readily extends both the PCHA and the FW algorithm to the objective in Equation 69. Specifically, to update $\mathbf{A}$ we can use the simply use the gradient shown in Equation 62. To update $\mathbf{B}$ and $\boldsymbol{\alpha}$ we need to derive their gradient with respect to Equation 69. Using $\breve{\mathbf{A}} = \mathbf{WA}$ and $\breve{\mathbf{X}} = \mathbf{W}\tilde{\mathbf{X}}$, we get for $\mathbf{B}$

$$\begin{aligned} \mathbf{G}^{(B)} &= \nabla_{\mathbf{B}}\,\mathrm{RSS} \\ &= 2\left[\mathrm{diag}(\boldsymbol{\alpha})\breve{\mathbf{A}}^T\breve{\mathbf{A}}\,\mathrm{diag}(\boldsymbol{\alpha})\mathbf{B}\tilde{\mathbf{X}}\tilde{\mathbf{X}}^T - \mathrm{diag}(\boldsymbol{\alpha})\breve{\mathbf{A}}^T\breve{\mathbf{X}}\tilde{\mathbf{X}}^T\right] \end{aligned} \tag{70}$$

as derived in Equation 140. For $\boldsymbol{\alpha}$ we have

$$\begin{aligned} \mathbf{G}^{(\alpha)} &= \nabla_{\boldsymbol{\alpha}}\,\mathrm{RSS} \\ &= 2\left[\breve{\mathbf{A}}^T\breve{\mathbf{A}}\,\mathrm{diag}(\boldsymbol{\alpha})\mathbf{B}\tilde{\mathbf{X}}\tilde{\mathbf{X}}^T\mathbf{B}^T - \breve{\mathbf{A}}^T\breve{\mathbf{X}}\tilde{\mathbf{X}}^T\mathbf{B}^T\right] \end{aligned} \tag{71}$$

as derived in Equation 141. The corresponding algorithm is outlined in Algorithm 12

**Algorithm 12** Prototypical Algorithm using Coresets and Relaxation of Archetypal Constraints

**Input:** Data matrix $\mathbf{X} \in \mathbb{R}^{N \times D}$, Subsampled data matrix $\tilde{\mathbf{X}} \in \mathbb{R}^{\tilde{N} \times D}$, coreset weights $\mathbf{W} \in \mathbb{R}_+^{\tilde{N} \times \tilde{N}}$ number of archetypes $K$, max iterations $T$
**Initialize:** Archetypes $\mathbf{Z} \in \mathbb{R}^{K \times D}$, $\boldsymbol{\alpha} = \mathbf{1}_K$
**for** $t = 1$ to $T$ **do**
  Compute optimal weights $\mathbf{A} \in \mathbb{R}^{N \times K}$:

$$\mathbf{A} \leftarrow \underset{\mathbf{A} \in F(\tilde{N}, K)}{\arg\min} \|\tilde{\mathbf{X}} - \mathbf{A}\,\mathrm{diag}(\boldsymbol{\alpha})\mathbf{B}\tilde{\mathbf{X}}\|_F^2 \tag{72}$$

  Compute optimal weights $\mathbf{B} \in \mathbb{R}^{K \times N}$:

$$\mathbf{B} \leftarrow \underset{\mathbf{B} \in F(K, \tilde{N})}{\arg\min} \|\mathbf{W}\tilde{\mathbf{X}} - \mathbf{W}\mathbf{A}\,\mathrm{diag}(\boldsymbol{\alpha})\mathbf{B}\tilde{\mathbf{X}}\|_F^2 \tag{73}$$

  Compute optimal $\boldsymbol{\alpha} \in [1-\delta, 1+\delta]^K$:

$$\boldsymbol{\alpha} \leftarrow \underset{\boldsymbol{\alpha} \in [1-\delta, 1+\delta]^K}{\arg\min} \|\mathbf{W}\tilde{\mathbf{X}} - \mathbf{W}\mathbf{A}\,\mathrm{diag}(\boldsymbol{\alpha})\mathbf{B}\tilde{\mathbf{X}}\|_F^2 \tag{74}$$

  Update archetypes:

$$\mathbf{Z} \leftarrow \mathrm{diag}(\boldsymbol{\alpha})\mathbf{B}\tilde{\mathbf{X}} \tag{75}$$

  **if** convergence criterion is met **then**
    **break**
  **end if**
**end for**
Recompute $\mathbf{A} \in F(N, K)$ on full dataset

$$\mathbf{A} \leftarrow \underset{\mathbf{A} \in F(N, K)}{\arg\min} \|\mathbf{X} - \mathbf{A}\mathbf{Z}\|_F^2 \tag{76}$$

**Return:** $\mathbf{A}, \mathbf{B}, \mathbf{Z}$

#### 2.8 Simulating Archetypes

##### 2.8.1 Archetype Generation

Let $K$ be the number of archetypes and $D$ the dimensionality of the feature space. We first generate a set of $M$ candidate points $C = \{c_1, \ldots, c_M\} \subset \mathbb{R}^D$, sampled independently from a standard multivariate normal

$$\mathbf{c}_i \sim \mathcal{N}(\mathbf{0}, \mathbf{I}_D) \tag{77}$$

We compute the convex hull $\mathrm{conv}(C)$ of the candidate set and extract its vertices:

$$\mathcal{V} = \mathrm{vertices}(\mathrm{conv}(C)) \subseteq C. \tag{78}$$

Thereby, we make sure that we will not have any redundant archetypes. Note however that the procedure is computationally tractable only for modest dimensions $D$; in high-dimensional spaces the convex-hull construction incurs a worst-case complexity of $\mathcal{O}\big(M^{\lfloor D/2 \rfloor}\big)$, making the overall algorithm prohibitively expensive.

If $|\mathcal{V}| \geq K$, we select a subset $Z = \{\mathbf{z}_1, \ldots, \mathbf{z}_K\} \subset \mathcal{V}$ by iteratively selecting points that maximize the minimum Euclidean distance to any of the points already selected. I.e. let $\mathcal{S} = \{\mathbf{v}_1, ..., \mathbf{v}_{t-1}\}$ be the set of points that have already been selected, then the next point is sampled according to:

$$\mathbf{v}_t = \arg\max_{\mathbf{v}\in\mathcal{V}} \min_{\mathbf{v}'\in\mathcal{S}} \|\mathbf{v} - \mathbf{v}'\|_2^2 \tag{79}$$

If $|\mathcal{V}| < K$, additional candidates are generated, and the process is repeated up to a maximum number of attempts.

###### 2.8.2 Coefficient Matrix Sampling

Given $N$ desired samples, we generate a matrix $\mathbf{A} \in \mathbb{R}^{N\times K}$ of mixing coefficients, where each row $\mathbf{a}_n = [a_{n1}, \ldots, a_{nK}]^T$ is sampled from a Dirichlet distribution

$$\mathbf{a}_n \sim \text{Dirichlet}(\mathbf{1}_K) \tag{80}$$

ensuring that each $\mathbf{a}_n \in \Delta_{K-1}$.

###### 2.8.3 Data Generation

Let $\mathbf{Z} \in \mathbb{R}^{K\times D}$ be the matrix whose rows are the selected archetypes. The synthetic data matrix $\mathbf{X} \in \mathbb{R}^{N\times D}$ is constructed via convex combinations of the archetypes:

$$\mathbf{X} = \mathbf{A}\mathbf{Z}. \tag{81}$$

Optionally, zero-mean isotropic Gaussian noise can be added to simulate measurement noise:

$$x_{nd} \leftarrow x_{nd} + \varepsilon_{nd}, \quad \varepsilon_{nd} \sim \mathcal{N}(0, \sigma^2), \tag{82}$$

where $\sigma$ is a user-defined noise standard deviation.

##### 2.9 Implementation Details

###### 2.9.1 Centering & Scaling

By default we center the data matrix $\mathbf{X}$, and globally scale $\mathbf{X}$ by

$$\lambda = \frac{\|X\|_F}{ND}. \tag{83}$$

This stabilizes the optimization, while translation and scaling invariance of the objective (see Section 2.2.1 and Section 2.2.2) ensure that the optimal $\mathbf{A}, \mathbf{B}$ are unaffected by this preprocessing.

###### 2.9.2 Convergence

Regardless of the objective and optimization algorithm, we use the moving average of the relative change in RSS, defined at iteration $t > 0$, as:

$$c_t = \frac{1}{\min(t, 20)} \sum_{\tau=1}^{\min(t,20)} \frac{\text{RSS}_{t-\tau} - \text{RSS}_{t-(\tau+1)}}{\text{RSS}_{t-(\tau+1)}} \tag{84}$$

We terminate the optimization if $c_t \geq 0$ or $|c_t| < 10^{-4}$.

###### 2.9.3 Multiple Restarts

By default, the `compute_archetypes` function runs the optimization five times with different random seeds, retaining only the result with the lowest RSS.

### 3 Number of Archetypes

There is no universally optimal method for selecting the number of archetypes (Alcacer et al. 2025). Here, we provide the following heuristics to guide the decision.

#### 3.1 Variance Explained

For a given number of archetypes, the variance explained is computed as

$$\begin{aligned} R^2 &= \frac{\text{TSS} - \text{RSS}}{\text{TSS}} \\ &= 1 - \frac{\text{RSS}}{\text{TSS}} \\ &= 1 - \frac{\|\mathbf{X} - \mathbf{ABX}\|_F^2}{\|\mathbf{X} - \mathbf{1}_N \bar{\mathbf{x}}^T\|_F^2} \end{aligned} \tag{85}$$

where $\mathbf{A}, \mathbf{B}$ denote the optimized coefficient matrices.

An "elbow" in this curve indicates that adding another archetype does not capture substantially more variance (Cutler and Breiman 1994; Eugster and Leisch 2009; Mørup and Hansen 2012).

#### 3.2 Information-Theoretic Criterion

Suleman (2017c) adapted the information-theoretic criterion from Suleman (2017a) for archetypal analysis, defining it as

$$v_{AA} = \log\left(\frac{1}{ND}\left\|\mathbf{X} - \hat{\mathbf{X}}\right\|_F^2\right) + 2\frac{K_a + K_b + 1}{N\,\text{tr}\left(\Sigma_{\hat{\mathbf{X}}}\Sigma_{\mathbf{X}}^{-1}\right)} \tag{86}$$

where $\hat{\mathbf{X}} = \mathbf{ABX}$ is the reconstructed data matrix; $K_a = N(K-1)$ (denoted $K_\mu$ in (Suleman 2017c)) is the number of parameters in $\mathbf{A}$; and $K_b = K(N-1)$ (denoted $K_\beta$ in (Suleman 2017c)) is the number of parameters in $\mathbf{B}$. The matrices $\Sigma_{\hat{\mathbf{X}}}$ and $\Sigma_{\mathbf{X}}$ denote the empirical covariances of $\hat{\mathbf{X}}$ and $\mathbf{X}$, respectively.

This criterion measures the goodness of fit by balancing model complexity and reconstruction accuracy: a lower value indicates a more favorable trade-off between the number of archetypes and the variance explained.

#### 3.3 Bootstrapping

Following the approach in (Hart et al. 2015; Korem et al. 2015), we assess the stability of the inferred archetypes using a bootstrap-based method.

For a given number of archetypes $K$, we first compute the optimal archetypes $\{\mathbf{z}_1, ..., \mathbf{z}_K\}$ on the full dataset.

Then, for each bootstrap sample $t = 1, ..., T$, we compute the corresponding optimal archetypes $\{\mathbf{z}_1^{(t)}, ..., \mathbf{z}_K^{(t)}\}$. These are matched to the original archetypes by solving the linear assignment problem:

$$\sigma^{(t)} = \arg\min_{\sigma \in S_K} \sum_{k=1}^{K} c\left(\mathbf{z}_k, \mathbf{z}_{\sigma(k)}^{(t)}\right), \tag{87}$$

where $S_K$ is the set of all permutations of $\{1, ..., K\}$, and the cost function $c$ is defined as the Euclidean distance, i.e.,

$$c(\mathbf{z}_k, \mathbf{z}_{k'}) = \|\mathbf{z}_k - \mathbf{z}_{k'}\|_2 \,. \tag{88}$$

We then compute the boostrap variance per archetype $k$ as

$$v_k = \frac{1}{TD} \sum_{t=1}^{T} \left\| \mathbf{z}_k^{(t)} - \bar{\mathbf{z}}_k \right\|_2^2, \tag{89}$$

where the mean archetype is defined as $\bar{\mathbf{z}}_k = \frac{1}{T} \sum_{t=1}^{T} \mathbf{z}_k^{(t)}$.

The variance $v_k$ serves as a measure of the stability of archetype $k$: a low variance indicates a robust and well-defined archetype, whereas a high variance suggests sensitivity to resampling.

### 4 Archetype Characterization

To characterize each archetype $k$, we define a smooth weighting over all cells $n$ using a squared exponential kernel:

$$w_{kn} = \frac{\exp\left[-\frac{\|\mathbf{z}_k - \mathbf{x}_n\|_2^2}{2\ell^2}\right]}{\sum_{n'=1}^{N} \exp\left[-\frac{\|\mathbf{z}_k - \mathbf{x}_{n'}\|_2^2}{2\ell^2}\right]}, \quad \mathbf{W} \in [0, 1]^{K \times N} \tag{90}$$

Thereby, cells that are close to an archetype get a large weight for that archetype.

By default, the length scale $\ell$ is determined automatically as

$$\ell = \frac{1}{2} \operatorname{median}\left(\{\|\mathbf{z}_k - \bar{\mathbf{x}}\|_2 \mid k \in \{1, ..., K\}\}\right) \tag{91}$$

where $\bar{\mathbf{x}} = \frac{1}{N} \sum_{n=1}^{N} \mathbf{x}_n$ denotes the data mean.

Note that the weight computation also implies that for each archetype $k$ we have

$$\mathbf{w}_k \geq 0 \text{ and } \sum_{n=1}^{N} w_{kn} = 1. \tag{92}$$

Multiplying the weight matrix $\mathbf{W} \in \mathbb{R}^{K \times N}$ with the original feature matrix $\mathbf{Y}^{(\mathbf{X})} \in \mathbb{R}^{N \times G}$ (where $G$ denotes the number of original features) yields a feature profile for each archetype.

$$\mathbf{Y}^{(\mathbf{Z})} = \mathbf{W}\mathbf{Y}^{(\mathbf{X})} \in \mathbb{R}^{K \times G} \tag{93}$$

For downstream analysis, we multiply the weight matrix with the z-scored, log1p-normalized expression matrix $\mathbf{Y}^{(\mathbf{X})}_{\text{z-scores}}$ (where z-scores are computed across cells). This highlights genes and biological processes that distinguish the archetypes, since uniformly expressed genes exhibit low z-scores, while cell state-specific genes display high z-scores.

Similarly, to associate continuous covariates (e.g. age, pseudotime) with archetypes, we multiply the weight matrix with the covariate vector.

For categorical covariates, we first one-hot encode the covariate yielding some matrix $\mathbf{H} \in \{0,1\}^{N \times C}$ where $C$ is the number of categories. To account for different number of samples per category, we normalize each column (i.e. category) $c$ to sum to one

$$\tilde{h}_{nc} = \frac{h_{nc}}{\sum_{n'=1}^{N} h_{n'c}}, \quad \tilde{\mathbf{H}} \in [0,1]^{N \times C} \tag{94}$$

This normalization ensures that archetype scores are not biased by class size. Then multiplying the weight matrix $\mathbf{W}$ with $\tilde{\mathbf{H}}$ yields enrichment scores per archetype per category.

$$\mathbf{S} = \mathbf{W}\tilde{\mathbf{H}} \in \mathbb{R}_{+}^{K \times C} \tag{95}$$

#### 4.1 Enrichment Analysis

To perform enrichment analysis we use decoupler-py (Badia-i-Mompel et al. 2022). By default, we use the univariate linear model (ULM) implemented in the package. Briefly, given some gene set with weights $\mathbf{w} = [w_1, ..., w_G]^T$, the ULM method models the expression profile of archetype $k$, denoted $\mathbf{y}_k^{(Z)}$, as

$$\mathbf{y}_k^{(Z)} = \beta_0 + \beta_1 \mathbf{w} + \epsilon_k. \tag{96}$$

Intuitively, if genes with higher weights tend to be more highly expressed in archetype $k$, the coefficient $\beta_1$ will be large. Conversely, if the weights are unrelated to expression, $\beta_1$ will be close to zero.

For unweighted gene sets, such as the Hallmark gene sets (Subramanian et al. 2005), weights are binary—set to one if a gene is in the set and zero otherwise.

The enrichment score is then calculated as the t-value of $\beta_1$

$$t_{\beta_1} = \frac{\hat{\beta}_1}{\text{SE}(\hat{\beta}_1)} \tag{97}$$

Note that the modular implementation of ParTIpy facilitates using any other enrichment analysis software.

#### 4.2 Spatial Mapping

Consider the setting where we have a *dissociated* single-cell dataset with log1p-normalized gene-expression matrix $\mathbf{Y} \in \mathbb{R}^{N \times G}$ and corresponding low-dimensional embedding $\mathbf{X} \in \mathbb{R}^{N \times D}$. In the same biological context, we also have a spatial single-cell dataset with log1p-normalized expression matrix $\mathbf{Y}' \in \mathbb{R}^{N' \times G'}$, where typically $G' < G$. Because the dissociated dataset captures more genes and therefore more information, we infer the archetypes on $\mathbf{X}$, yielding the characteristic expression profiles $\mathbf{Y}^{(\mathbf{Z})} \in \mathbb{R}^{K \times G}$ as described above. To relate the spatial cells to these archetypes, we restrict $\mathbf{Y}^{(\mathbf{Z})}$ to the intersection of genes between the two datasets (denoting the size of the intersection with $\mathbf{G}''$), obtaining $\mathbf{Y}^{(\mathbf{Z})}_{\mathcal{G}} \in \mathbb{R}^{K \times G''}$. We then apply decoupler-py's univariate linear model (ULM) (Badia-i-Mompel et al. 2022) to each profile $\mathbf{y}^{(\mathbf{Z})}_{k,\mathcal{G}}$ to compute enrichment scores which quantify how strongly each cell $n'$ in the spatial dataset is associated with each archetype $k$.

#### 4.3 Archetype Crosstalk Networks

The observed distribution of cells between archetypes can be understood as functional *division of labor* between the task that each archetype is specialized in. Ligand-receptor (LR) signaling (e.g. lateral inhibition) is one potential mechanism that establishes and maintains this pattern of specialization (Adler et al. 2023). To map the possible lines of communication between archetypes, we combine their characteristic expression profiles with LR databases (Dimitrov et al. 2022; Dimitrov et al. 2024), following the workflow adapted from Adler et al. (2023).

**Step 1: Identify archetype-enriched genes.** For each archetype $k$ and gene $g$ we compute a specificity score

$$s_{kg} = y^{(\mathbf{Z})}_{kg} - \max_{k' \neq k} y^{(\mathbf{Z})}_{k'g}, \tag{98}$$

where $\mathbf{Y}^{(\mathbf{Z})} \in \mathbb{R}^{K \times G}$ is the matrix of characteristic expression profiles inferred using the z-scored log1p single-cell expression profiles. Genes with $s_{kg} > \tau$ (default $\tau = 0.1$) are deemed *enriched* in archetype $k$.

**Step 2: Filter to archetype-specific LR pairs.** Let $\mathcal{D} = \{(l, r)\}$ be a curated set of cognate LR pairs (Dimitrov et al. 2022; Dimitrov et al. 2024). For every ordered pair of archetype $(k, k')$, we retain only those pairs whose ligand is enriched in $k$ and whose receptor is enriched in $k'$:

$$\mathcal{D}_{k \to k'} = \{(l, r) \in \mathcal{D} : l \text{ enriched in } k,\ r \text{ enriched in } k'\}. \tag{99}$$

**Step 3: Build archetype crosstalk network** For every ordered pair $(k, k')$ with $|\mathcal{D}_{k \to k'}| > 0$ we insert a directed edge $k \to k'$ and assign it the weight $w_{k \to k'} = |\mathcal{D}_{k \to k'}|$.

The resulting weighted graph summarizes the potential signaling axes that may coordinate the division of labor among archetypes.

### 5 Benchmarks

#### 5.1 Benchmarking Initialization & Optimization Algorithms

To determine the default initialization and optimization algorithm, we benchmarked all three initialization strategies in combination with two optimization methods (PCHA and FW). RNNLS was excluded due to scalability limitations. The benchmark was conducted on the datasets and cell types listed in Supplementary Table 1. For each combination of initialization and optimization, we ran the optimization procedure with 20 different random seeds per cell type, recording both the variance explained and the elapsed time. Results for each study are shown in Supplementary Figure S1 (panels C-E). To aggregate results across cell types, we z-scored the variance explained and elapsed time within each cell type (i.e., subtracting the mean and dividing by the standard deviation), and then computed the mean and standard deviation of these z-scores across all experiments. The summary is presented in Supplementary Figure S1B.

##### 5.2 Benchmarking Coresets

To derive recommendations for optimal coreset sizes, we evaluated performance across a range of coreset size fractions [1.0, 0.64, 0.32, 0.16, 0.08, 0.04, 0.02, 0.01, 0.005]. For each size, we performed 20 independent optimization runs (using different random seeds), recording both the variance explained and the elapsed time. The same datasets and cell types as in Section 5.1 were used (see Supplementary Table 1). Study-specific results are presented in Supplementary Figure 3, panels C-E. For each cell type, we fitted a Generalized Additive Model (GAM) to model the relationship between coreset size and variance explained. The optimal coreset size was then defined as the smallest size for which the predicted variance explained remained within 5% of that obtained using the full dataset. These optimal coreset sizes formed the basis for the summary statistics shown in Supplementary Figure 3, panels A-B.

#### 6 Data Analysis

##### 6.1 Hepatocytes

We applied our framework to the hepatocyte dataset from Ben-Moshe et al. (2022), which comprises 1,999 cells and 8,354 genes. Specifically, we restricted our analysis to hepatocytes collected at time point zero, prior to the acetaminophen (APAP)-induced injury and subsequent regenerative response studied by Ben-Moshe et al. (2022). Preprocessing was conducted using scanpy (Wolf, Angerer, and Theis 2018), following the steps outlined below. First, total counts per cell were scaled to the median library size across all cells, followed by log1p transformation of the scaled counts. Principal component analysis (PCA) was then performed on the transformed expression matrix, using as input the subset of highly variable genes identified with the default algorithm in scanpy. The first three principal components were retained for downstream analysis.

Archetypal analysis was conducted using the default parameters, specifically Archetypal++ initialization and PCHA optimization. Based on the model selection criteria shown in Supplementary Figure 4A–C, we selected $K = 4$ archetypes. Characteristic gene expression profiles for each archetype were computed as described in Section 4, using z-scored log1p-normalized expression values.

To annotate the functional specialization of each archetype, we performed pathway enrichment analysis using the Reactome database (Gillespie et al. 2022) in combination with the univariate linear model (ULM) implemented in decoupler-py (Badia-i-Mompel et al. 2022). Spatial mapping was performed by projecting archetype gene expression profiles onto a spatial transcriptomics dataset using univariate linear modeling as described in Section 4.2. An archetype crosstalk network was inferred using the mouse consensus resource from LIANA+ (Dimitrov et al. 2024) following the procedure in Section 4.3

#### 7 References

Adler, Miri et al. (May 2023). "Emergence of Division of Labor in Tissues through Cell Interactions and Spatial Cues". In: *Cell Reports* 42.5, p. 112412. ISSN: 22111247. DOI: 10.1016/j.celrep.2023.112412. PMID: 37086403. URL: https://linkinghub.elsevier.com/retrieve/pii/S2211124723004230 (visited on 12/07/2024).

Alcacer, Aleix et al. (Apr. 16, 2025). *A Survey on Archetypal Analysis*. DOI: 10.48550/arXiv.2504.12392. arXiv: 2504.12392 [stat]. URL: http://arxiv.org/abs/2504.12392 (visited on 04/22/2025).

Bachem, Olivier, Mario Lucic, and Andreas Krause (July 19, 2018). "Scalable k -Means Clustering via Lightweight Coresets". In: *Proceedings of the 24th ACM SIGKDD International Conference on Knowledge Discovery & Data Mining*. KDD '18. New York, NY, USA: Association for Computing

Machinery, pp. 1119–1127. ISBN: 978-1-4503-5552-0. DOI: 10.1145/3219819.3219973. URL: https://dl.acm.org/doi/10.1145/3219819.3219973 (visited on 05/04/2025).

Badia-i-Mompel, Pau et al. (Jan. 2022). "decoupleR: Ensemble of Computational Methods to Infer Biological Activities from Omics Data". In: *Bioinformatics Advances* 2.1. Ed. by Marieke Lydia Kuijjer, vbac016. ISSN: 2635-0041. DOI: 10.1093/bioadv/vbac016. (Visited on 12/07/2024).

Bauckhage, Christian et al. (2015). "Archetypal Analysis as an Autoencoder". In: Workshop "New Challenges in Neural Computation" (NC2) 2015. URL: https://www.ml.informatik.tu-darmstadt.de/papers/autoencode2015nc2.pdf (visited on 02/10/2025).

Ben-Moshe, Shani et al. (June 2022). "The Spatiotemporal Program of Zonal Liver Regeneration Following Acute Injury". In: *Cell Stem Cell* 29.6, 973–989.e10. ISSN: 19345909. DOI: 10.1016/j.stem.2022.04.008. PMID: 35659879. URL: https://linkinghub.elsevier.com/retrieve/pii/S1934590922001618 (visited on 12/07/2024).

Clarkson, Kenneth L. (Sept. 3, 2010). "Coresets, Sparse Greedy Approximation, and the Frank-Wolfe Algorithm". In: *ACM Trans. Algorithms* 6.4, 63:1–63:30. ISSN: 1549-6325. DOI: 10.1145/1824777.1824783. URL: https://doi.org/10.1145/1824777.1824783 (visited on 02/11/2025).

Cutler, Adele and Leo Breiman (1994). "Archetypal Analysis". In: *Technometrics : a journal of statistics for the physical, chemical, and engineering sciences* 36.4, pp. 338–347. ISSN: 0040-1706. DOI: 10.1080/00401706.1994.10485840.

Dimitrov, Daniel et al. (June 2022). "Comparison of Methods and Resources for Cell-Cell Communication Inference from Single-Cell RNA-Seq Data". In: *Nature Communications* 13.1, p. 3224. ISSN: 2041-1723. DOI: 10.1038/s41467-022-30755-0. (Visited on 12/07/2024).

Dimitrov, Daniel et al. (Sept. 2024). "LIANA+ Provides an All-in-One Framework for Cell–Cell Communication Inference". In: *Nature Cell Biology* 26.9, pp. 1613–1622. ISSN: 1465-7392, 1476-4679. DOI: 10.1038/s41556-024-01469-w. (Visited on 12/07/2024).

Eggert, J. and E. Korner (July 2004). "Sparse Coding and NMF". In: *2004 IEEE International Joint Conference on Neural Networks (IEEE Cat. No.04CH37541).* Vol. 4, 2529–2533 vol.4. DOI: 10.1109/IJCNN.2004.1381036. URL: https://ieeexplore.ieee.org/document/1381036 (visited on 03/30/2025).

Eugster, Manuel J. A. and Friedrich Leisch (2009). "From Spider-Man to Hero - Archetypal Analysis in *R*". In: *Journal of Statistical Software* 30.8. ISSN: 1548-7660. DOI: 10.18637/jss.v030.i08. URL: http://www.jstatsoft.org/v30/i08/ (visited on 12/07/2024).

Feldman, Dan (Nov. 2020). "Introduction to Core-Sets: An Updated Survey". In: arXiv:2011.09384. DOI: 10.48550/arXiv.2011.09384. arXiv: 2011.09384 [cs]. (Visited on 07/02/2025).

Frank, Marguerite and Philip Wolfe (Mar. 1956). "An Algorithm for Quadratic Programming". In: *Naval Research Logistics Quarterly* 3.1-2, pp. 95–110. ISSN: 0028-1441, 1931-9193. DOI: 10.1002/nav.3800030109. (Visited on 06/29/2025).

Gillespie, Marc et al. (Jan. 2022). "The Reactome Pathway Knowledgebase 2022". In: *Nucleic Acids Research* 50.D1, pp. D687–D692. ISSN: 0305-1048, 1362-4962. DOI: 10.1093/nar/gkab1028. (Visited on 12/07/2024).

Hart, Yuval et al. (Mar. 2015). "Inferring Biological Tasks Using Pareto Analysis of High-Dimensional Data". In: *Nature Methods* 12.3, pp. 233–235. ISSN: 1548-7091, 1548-7105. DOI: 10.1038/nmeth.3254. PMID: 25622107. URL: https://www.nature.com/articles/nmeth.3254 (visited on 12/07/2024).

Hochbaum, Dorit S. and David B. Shmoys (May 1985). "A Best Possible Heuristic for the K-Center Problem". In: *Mathematics of Operations Research* 10.2, pp. 180–184. ISSN: 0364-765X. DOI: 10.1287/moor.10.2.180. (Visited on 06/08/2025).

Jaggi, Martin (Feb. 13, 2013). "Revisiting Frank-Wolfe: Projection-Free Sparse Convex Optimization". In: *Proceedings of the 30th International Conference on Machine Learning.* International Conference on Machine Learning. PMLR, pp. 427–435. URL: https://proceedings.mlr.press/v28/jaggi13.html (visited on 02/08/2025).

Korem, Yael et al. (July 10, 2015). "Geometry of the Gene Expression Space of Individual Cells". In: *PLOS Computational Biology* 11.7. Ed. by Lilia M. Iakoucheva, e1004224. ISSN: 1553-7358. DOI: 10.1371/journal.pcbi.1004224. PMID: 26161936. URL: https://dx.plos.org/10.1371/journal.pcbi.1004224 (visited on 12/07/2024).

Krohne, Laerke Gebser et al. (Aug. 2019). "Classification of Social Anhedonia Using Temporal and Spatial Network Features from a Social Cognition fMRI Task". In: *Human Brain Mapping* 40.17, pp. 4965–4981. ISSN: 1065-9471. DOI: 10.1002/hbm.24751. (Visited on 06/08/2025).

Lin, Chih-Jen (Oct. 2007). "Projected Gradient Methods for Nonnegative Matrix Factorization". In: *Neural Computation* 19.10, pp. 2756–2779. ISSN: 0899-7667. DOI: 10.1162/neco.2007.19.10.2756. URL: https://ieeexplore.ieee.org/document/6795860 (visited on 03/30/2025).

Mair, Sebastian and Ulf Brefeld (2019). "Coresets for Archetypal Analysis". In: *Advances in Neural Information Processing Systems*. Vol. 32. Curran Associates, Inc. URL: https://papers.nips.cc/paper_files/paper/2019/hash/7f278ad602c7f47aa76d1bfc90f20263-Abstract.html (visited on 02/09/2025).

Mair, Sebastian and Jens Sjölund (May 13, 2024). *Archetypal Analysis++: Rethinking the Initialization Strategy*. DOI: 10.48550/arXiv.2301.13748. arXiv: 2301.13748 [cs]. URL: http://arxiv.org/abs/2301.13748 (visited on 12/07/2024). Pre-published.

Minc, Henryk (1988). *Nonnegative Matrices*. 2. print. Wiley-Interscience Series in Discrete Mathematics and Optimization. New York: Wiley. 206 pp. ISBN: 978-0-471-83966-8.

Mørup, Morten and Lars Kai Hansen (Mar. 2012). "Archetypal Analysis for Machine Learning and Data Mining". In: *Neurocomputing* 80, pp. 54–63. ISSN: 09252312. DOI: 10.1016/j.neucom.2011.06.033. URL: https://linkinghub.elsevier.com/retrieve/pii/S0925231211006060 (visited on 12/07/2024).

Olsen, Anders S. et al. (July 2022). "Combining Electro- and Magnetoencephalography Data Using Directional Archetypal Analysis". In: *Frontiers in Neuroscience* 16. ISSN: 1662-453X. DOI: 10.3389/fnins.2022.911034. (Visited on 06/08/2025).

Petersen, Kaare Brandt and Michael Syskind Pedersen (2012). *The Matrix Cookbook*. Technical University of Denmark. URL: https://www.math.uwaterloo.ca/~hwolkowi/matrixcookbook.pdf.

Subramanian, Aravind et al. (Oct. 2005). "Gene Set Enrichment Analysis: A Knowledge-Based Approach for Interpreting Genome-Wide Expression Profiles". In: *Proceedings of the National Academy of Sciences* 102.43, pp. 15545–15550. ISSN: 0027-8424, 1091-6490. DOI: 10.1073/pnas.0506580102. (Visited on 12/07/2024).

Suleman, Abdul (Mar. 2017a). "Measuring the Congruence of Fuzzy Partitions in Fuzzy $C$-Means Clustering". In: *Applied Soft Computing* 52, pp. 1285–1295. ISSN: 1568-4946. DOI: 10.1016/j.asoc.2016.06.037. (Visited on 06/09/2025).

— (Dec. 2017b). "On Ill-Conceived Initialization in Archetypal Analysis". In: *Advances in Data Analysis and Classification* 11.4, pp. 785–808. ISSN: 1862-5355. DOI: 10.1007/s11634-017-0303-0. (Visited on 06/08/2025).

— (July 2017c). "Validation of Archetypal Analysis". In: *2017 IEEE International Conference on Fuzzy Systems (FUZZ-IEEE)*. 2017 IEEE International Conference on Fuzzy Systems (FUZZ-IEEE), pp. 1–6. DOI: 10.1109/FUZZ-IEEE.2017.8015385. URL: https://ieeexplore.ieee.org/document/8015385 (visited on 04/23/2025).

Wolf, F. Alexander, Philipp Angerer, and Fabian J. Theis (Dec. 2018). "SCANPY: Large-Scale Single-Cell Gene Expression Data Analysis". In: *Genome Biology* 19.1, p. 15. ISSN: 1474-760X. DOI: 10.1186/s13059-017-1382-0. (Visited on 12/07/2024).

### 8 Appendix

#### 8.1 Proof for Translation Invariance

Let $\mathbf{v} \in \mathbb{R}^D$, and let $\tilde{\mathbf{X}} = \mathbf{X} + \mathbf{1}_N \mathbf{v}^T$ be the translated matrix. Then for any feasible $\mathbf{A}, \mathbf{B}$

$$\begin{aligned} \tilde{\mathbf{X}} - \mathbf{A}\mathbf{B}\tilde{\mathbf{X}} &= \left(\mathbf{X} + \mathbf{1}_N \mathbf{v}^T\right) - \mathbf{A}\mathbf{B}\left(\mathbf{X} + \mathbf{1}_N \mathbf{v}^T\right) \\ &= \mathbf{X} + \mathbf{1}_N \mathbf{v}^T - \mathbf{A}\mathbf{B}\mathbf{X} - \mathbf{A}\mathbf{B}\mathbf{1}_N \mathbf{v}^T \end{aligned} \tag{100}$$

Since $\mathbf{B}\mathbf{1}_N = \mathbf{1}_K$ and $\mathbf{A}\mathbf{1}_K = \mathbf{1}_N$, this simplifies to

$$\begin{aligned} \tilde{\mathbf{X}} - \mathbf{A}\mathbf{B}\tilde{\mathbf{X}} &= \mathbf{X} + \mathbf{1}_N \mathbf{v}^T - \mathbf{A}\mathbf{B}\mathbf{X} - \mathbf{1}_N \mathbf{v}^T \\ &= \mathbf{X} - \mathbf{A}\mathbf{B}\mathbf{X} \end{aligned} \tag{101}$$

Therefore, the reconstruction error remains unchanged, and the minimizers $\mathbf{A}, \mathbf{B}$ are invariant under such translations. Thus, the minimizers $\mathbf{A}, \mathbf{B}$ are invariant to centering the data.

#### 8.2 Proof for Scaling Invariance

Let $\lambda \neq 0$, and let $\tilde{\mathbf{X}} = \lambda \mathbf{X}$ be the scaled matrix. Then for any feasible $\mathbf{A}, \mathbf{B}$

$$\begin{aligned} \tilde{\mathbf{X}} - \mathbf{A}\mathbf{B}\tilde{\mathbf{X}} &= \lambda \mathbf{X} - \mathbf{A}\mathbf{B}\lambda \mathbf{X} \\ &= \lambda \left(\mathbf{X} - \mathbf{A}\mathbf{B}\mathbf{X}\right) \end{aligned} \tag{102}$$

Thus the objective for the scaled matrix is given by

$$\underset{\substack{\mathbf{A} \in F(N,K) \\ \mathbf{B} \in F(K,N)}}{\arg\min} \|\tilde{\mathbf{X}} - \mathbf{A}\mathbf{B}\tilde{\mathbf{X}}\|_F^2 = \underset{\substack{\mathbf{A} \in F(N,K) \\ \mathbf{B} \in F(K,N)}}{\arg\min} \lambda^2 \|\mathbf{X} - \mathbf{A}\mathbf{B}\mathbf{X}\|_F^2 \tag{103}$$

Since $\lambda \neq 0$, we have $\lambda^2 > 0$, and thus the objective is scaled by a positive constant. Multiplying the objective function by a positive scalar does not affect the location of its minimum, because the ordering of objective values is preserved. In particular, the first-order (stationarity) and second-order (convexity/curvature) necessary conditions for optimality remain unchanged under such scaling.

#### 8.3 Proof for Uniqueness up to Permutation

Assuming that for each archetype $k$, there exists one data point $n$ that is best reconstructed using only this archetype, i.e.

$$\forall k \in \{1, ..., K\} \exists n \in \{1, ..., N\} \; a_{nk} = 1 \tag{104}$$

and that for each archetype there exists one data point that is only used to define this archetype and not any other archetype, i.e.

$$\forall k \in \{1, ..., K\} \exists n \in \{1, ..., N\} \; b_{kn} > 0 \;\wedge\; b_{k'n} = 0 \forall k' \neq k \tag{105}$$

then the archetypes are identifiable up to permutation (i.e. the objective does not suffer from rotational ambiguity).

Formally, let $\mathbf{A} \in F(N, K)$ and $\mathbf{B} \in F(K, N)$ be feasible, and let the assumption in Equation 104 and Equation 105 hold. Consider any invertible $\mathbf{Q} \in \mathbb{R}^{K \times K}$ such that the transformed pair

$$\mathbf{A}' = \mathbf{A}\mathbf{Q}, \qquad \mathbf{B}' = \mathbf{Q}^{-1}\mathbf{B} \tag{106}$$

is also feasible, i.e. $\mathbf{A}' \in F(N, K)$ and $\mathbf{B}' \in F(K, N)$. Then $\mathbf{Q}$ must be a permutation matrix.

Condition 1 implies that $\mathbf{A}$ has at least $K$ rows that have only one non-zero value, and this non-zero value equals one.

Condition 2 implies that $\mathbf{B}$ has at least $K$ columns that have only one non-zero value.

Example: If we had 3 archetypes, the following weight matrices $\mathbf{A}$, $\mathbf{B}$ would satisfy these conditions

$$\mathbf{A} = \begin{bmatrix} \vdots & \vdots & \vdots \\ 1 & 0 & 0 \\ \vdots & \vdots & \vdots \\ 0 & 0 & 1 \\ \vdots & \vdots & \vdots \\ 0 & 1 & 0 \\ \vdots & \vdots & \vdots \end{bmatrix}, \quad \mathbf{B} = \begin{bmatrix} \cdots & 0 & \cdots & 0.4 & \cdots & 0 & \cdots \\ \cdots & 0 & \cdots & 0 & \cdots & 0.1 & \cdots \\ \cdots & 0.2 & \cdots & 0 & \cdots & 0 & \cdots \end{bmatrix} \tag{107}$$

Given the above assumptions, we can derive the following properties that $\mathbf{Q}$ must fulfill.

*First,* $\mathbf{Q} \geq 0$.

Proof: By the condition in Equation (104), we assume that for each $k \in \{1, ..., K\}$, $\mathbf{A}$ contains at least one row with a single non-zero entry which we denote by $n_k$. Now, $(\mathbf{A}\mathbf{Q})_{n_k,:} = \mathbf{e}_k^\top \mathbf{Q}$ is the $k$th row of $\mathbf{Q}$. Since we require that $\mathbf{A}\mathbf{Q} \geq 0$, every row of $\mathbf{Q}$ is entrywise nonnegative, and thus $\mathbf{Q}$ is nonnegative.

*Second,* $\mathbf{Q}^{-1} \geq 0$.

Proof: By the condition in Equation (105), we assume that for each $k \in \{1, ..., K\}$, $\mathbf{B}$ contains at least one column with a single non-zero entry which we denote by $n_k$. Then $(\mathbf{B}')_{:,n_k} = \mathbf{Q}^{-1}\mathbf{B}_{:,n_k} = \mathbf{Q}^{-1}_{:,n_k} b_{k,n_k}$. Since we know that $\mathbf{Q}^{-1}_{:,n_k} b_{k,n_k} \geq 0$ and $b_{k,n_k} > 0$, we have $\mathbf{Q}^{-1}_{:,n_k} \geq 0$ and thus $\mathbf{Q}^{-1} \geq 0$.

*Third,* $\mathbf{Q}$ *is a generalized permutation matrix.*

Since $\mathbf{Q} \geq 0$ and $\mathbf{Q}^{-1} \geq 0$, by Minc's Lemma 1.1 Minc (1988) there exist a diagonal $\mathbf{D} = \mathrm{diag}(d_1, \ldots, d_K)$ with $d_k > 0$ and a permutation matrix $\mathbf{P}$ such that $\mathbf{Q} = \mathbf{D}\mathbf{P}$.

*Fourth,* $\mathbf{Q}$ *is a permutation.*

Proof: Given that $\mathbf{P}\mathbf{1}_K = \mathbf{1}_K$, we have

$$\begin{aligned} &\mathbf{A}\mathbf{Q}\mathbf{1}_K = \mathbf{1}_N \\ \rightarrow &\mathbf{A}\mathbf{D}\mathbf{P}\mathbf{1}_K = \mathbf{1}_N \\ \rightarrow &\mathbf{A}\mathbf{D}\mathbf{1}_K = \mathbf{1}_N. \end{aligned} \tag{108}$$

Thus, for each $n \in 1, ..., N$ we require $\sum_{k=1}^{K} a_{nk} d_k = 1$. By the condition in Equation (104), we assume that for each $k \in \{1, ..., K\}$, $\mathbf{A}$ contains at least one row with a single non-zero entry. Thus to satisfy $\sum_{k=1}^{K} a_{nk} d_k = 1$ for these rows, $d_k = 1$. Thus $\mathbf{D} = \mathbf{I}_K$ and $\mathbf{Q} = \mathbf{D}\mathbf{P} = \mathbf{P}$.

#### 8.4 Proof for Convexity of Objective

If we measure the reconstruction error with the RSS, the objective in Eq. (4) is biconvex: it is convex in $\mathbf{A}$ when $\mathbf{B}$ is fixed and vice-versa. We prove convexity in $\mathbf{A}$; the argument for $\mathbf{B}$ is analogous.

Let $Z := \mathbf{BX} \in \mathbb{R}^{K\times D}$. With $\mathbf{B}$ fixed, Eq. (4) reduces to the quadratic programme

$$\hat{\mathbf{A}} = \underset{\mathbf{A}\in\mathbb{R}^{N\times K}}{\arg\min} \|\mathbf{X}-\mathbf{AZ}\|_F^2 \quad \text{s.t.} \quad \mathbf{A}\geq 0,\ \mathbf{A}\mathbf{1}_K = \mathbf{1}_N. \tag{109}$$

As shown in Section 8.5, the feasible set is convex. We now rewrite the objective in canonical quadratic form.

Defining the vectorizations

$$\begin{aligned} \operatorname{vec}(\mathbf{X}) &= \begin{bmatrix} x_{11} & \dots & x_{1D} & \dots & x_{N1} & \dots & x_{ND} \end{bmatrix}^T \in \mathbb{R}^{ND} \\ \operatorname{vec}(\mathbf{A}) &= \begin{bmatrix} a_{11} & \dots & a_{1K} & \dots & a_{N1} & \dots & a_{NK} \end{bmatrix}^T \in \mathbb{R}_+^{NK} \end{aligned} \tag{110}$$

and Kronecker stack $\mathbf{Z}_\otimes$

$$\mathbf{Z}_\otimes = \begin{bmatrix} \mathbf{Z}^T & 0 & 0 & 0 \\ 0 & \mathbf{Z}^T & 0 & 0 \\ & & \ddots & \\ 0 & 0 & & 0 \\ 0 & 0 & 0 & \mathbf{Z}^T \end{bmatrix} = \mathbf{I}_N \otimes \mathbf{Z}^T \in \mathbb{R}^{ND\times NK} \tag{111}$$

we can rewrite the squared Frobenius norm as squared $\ell_2$-norm

$$\begin{aligned} \|\mathbf{X}-\mathbf{AZ}\|_F^2 &= \left\|\operatorname{vec}(\mathbf{X}) - \mathbf{Z}_\otimes \operatorname{vec}(\mathbf{A})\right\|_2^2 \\ &= \left(\operatorname{vec}(\mathbf{X}) - \mathbf{Z}_\otimes \operatorname{vec}(\mathbf{A})\right)^T \left(\operatorname{vec}(\mathbf{X}) - \mathbf{Z}_\otimes \operatorname{vec}(\mathbf{A})\right) \\ &= \operatorname{vec}(\mathbf{X})^T \operatorname{vec}(\mathbf{X}) - 2\operatorname{vec}(\mathbf{X})^T \mathbf{Z}_\otimes \operatorname{vec}(\mathbf{A}) + \operatorname{vec}(\mathbf{A})^T \mathbf{Z}_\otimes^T \mathbf{Z}_\otimes \operatorname{vec}(\mathbf{A}). \end{aligned} \tag{112}$$

Thus, the optimization problem can be written in the canonical form of a quadratic program

$$\underset{\operatorname{vec}(\mathbf{A})\in\mathbb{R}^{NK}}{\arg\min} \frac{1}{2}\operatorname{vec}(\mathbf{A})^T \mathbf{Q} \operatorname{vec}(\mathbf{A}) + \mathbf{c}^T \operatorname{vec}(\mathbf{A}) \quad \text{s.t.} \quad \operatorname{vec}(\mathbf{A}) \geq 0,\ \mathbf{C}\operatorname{vec}(\mathbf{A}) = \mathbf{1}_N \tag{113}$$

with

$$\begin{aligned} \frac{1}{2}\mathbf{Q} &= \mathbf{Z}_\otimes^T \mathbf{Z}_\otimes = (\mathbf{I}_N \otimes \mathbf{Z}^T)^T (\mathbf{I}_N \otimes \mathbf{Z}^T) = (\mathbf{I}_N^T \otimes \mathbf{Z})(\mathbf{I}_N \otimes \mathbf{Z}^T) = \mathbf{I}_N^T \mathbf{I}_N \otimes \mathbf{Z}\mathbf{Z}^T = \mathbf{I}_N \otimes \mathbf{Z}\mathbf{Z}^T \\ \mathbf{c}^T &= -2\operatorname{vec}(\mathbf{X})^T \mathbf{Z}_\otimes \\ \mathbf{C} &= \mathbf{1}_K^T \otimes \mathbf{I}_N \end{aligned} \tag{114}$$

Because $\mathbf{Z}\mathbf{Z}^T \geq 0$, the Kronecker product $\mathbf{I}_N \otimes (\mathbf{Z}\mathbf{Z}^\top)$ is also positive semi-definite, and scaling by 2 preserves this property. Hence $\mathbf{Q} \geq 0$.

The objective in (109) is a convex quadratic, and the constraints are linear; therefore the $\mathbf{A}$-subproblem is a convex quadratic programme. Fixing $\mathbf{A}$ and repeating the same construction for $\mathbf{B}$ proves the corresponding convexity in $\mathbf{B}$.

#### 8.5 Proof for Convexity of Standard Simplex

Let $\mathbf{x}, \mathbf{y} \in \Delta_{K-1}$, then for all $0 \leq \lambda \leq 1$, it is true that $\mathbf{z} = \lambda\mathbf{x} + (1-\lambda)\mathbf{y} \in \Delta_{K-1}$

First we show that for all $k \in \{1, ..., K\}$, $z_k \geq 0$. Recall that $z_k = \lambda x_k + (1-\lambda)y_k$. Since $0 \leq \lambda \leq 1$, we have $1 - \lambda \geq 0$. Since $\mathbf{x}, \mathbf{y} \in \Delta_{K-1}$, we have $x_k \geq 0$ and $y_k \geq 0$. Thus $z_k = \lambda x_k + (1-\lambda)y_k \geq 0$.

Then, we show that the elements of $\mathbf{z}$ sum to one.

$$\begin{aligned}
\|\mathbf{z}\|_1 &= \sum_{k=1}^{K} z_k \\
&= \sum_{k=1}^{K} [\lambda x_k + (1-\lambda)y_k] \\
&= \lambda \underbrace{\sum_{k=1}^{K} x_k}_{=1} + (1-\lambda) \underbrace{\sum_{k=1}^{K} y_k}_{=1} \\
&= \lambda + 1 - \lambda \\
&= 1
\end{aligned}$$

#### 8.6 Proof for Compactness of Standard Simplex

The standard $(K-1)$-simplex $\Delta_{K-1}$

$$\Delta_{K-1} := \left\{ \mathbf{x} \in \mathbb{R}^K \mid \forall k \in \{1, ..., K\}, x_k \geq 0 \land \sum_{k=1}^{K} x_k = 1 \right\}. \tag{115}$$

is a subset of a finite-dimensional Euclidean space. By the *Heine-Borel theorem*, a subset of $\mathbb{R}^K$ is compact if and only if it is both *closed* and *bounded.*

**Step 1: Closedness.**

We can write $\Delta_{K-1}$ as the finite intersection

$$\Delta_{K-1} = \Big( \bigcap_{k=1}^{K} \underbrace{\{\mathbf{x} \in \mathbb{R}^K : x_k \geq 0\}}_{=:A_k} \Big) \cap \underbrace{\{\mathbf{x} \in \mathbb{R}^K : \sum_{k=1}^{K} x_k = 1\}}_{=:B}. \tag{116}$$

- *Each $A_k$ is closed.* Let $\pi_k : \mathbb{R}^K \to \mathbb{R}$, $\pi_k(\mathbf{x}) = x_k$. The coordinate projection $\pi_k$ is linear, hence continuous. Because $[0, \infty) \subset \mathbb{R}$ is closed, its pre-image

$$A_k = \pi_k^{-1}\big([0, \infty)\big) \tag{117}$$

  is closed. Geometrically each $A_k$ is a closed half-space.
- *$B$ is closed.* Define $g : \mathbb{R}^K \to \mathbb{R}$, $g(\mathbf{x}) = \sum_{k=1}^{K} x_k$. The set $B$ can be written as $g^{-1}(\{1\})$. Since $g$ is continuous and $\{1\}$ is closed in $\mathbb{R}$, $B$ is closed.

Because a *finite* intersection of closed sets is closed, it follows that $\Delta_{K-1}$ is closed.

**Step 2: Boundedness.**

For $\mathbf{x} \in \Delta_{K-1}$ we have $0 \le x_k \le 1$ and $\sum_{k=1}^{K} x_k = 1$. Since $x_k^2 \le x_k$ on $[0, 1]$,

$$\|\mathbf{x}\|_2^2 = \sum_{k=1}^{K} x_k^2 \le \sum_{k=1}^{K} x_k = 1, \quad \Longrightarrow \quad \|\mathbf{x}\|_2 \le 1. \tag{118}$$

Hence every element of $\Delta_{K-1}$ lies in the closed Euclidean ball $B_2(0, 1)$. Hence, the simplex is bounded.

**Step 3: Compactness.**

Since $\Delta_{K-1}$ is both closed (Step 1) and bounded (Step 2), the Heine-Borel theorem implies that $\Delta_{K-1}$ is compact in $\mathbb{R}^K$.

#### 8.7 Gradient of Vanilla Objective

To compute the gradient of the unconstrained objective with respect to $\mathbf{A}$ and $\mathbf{B}$, we first rewrite the residual sum of squares (Frobenius norm) in Equation (4) in terms of the trace

$$\begin{aligned} \text{RSS} &= \|\mathbf{X} - \mathbf{ABX}\|_F^2 \\ &= \operatorname{tr}\left((\mathbf{X} - \mathbf{ABX})^T (\mathbf{X} - \mathbf{ABX})\right) \\ &= \operatorname{tr}(\mathbf{X}^T\mathbf{X}) - \operatorname{tr}(\mathbf{X}^T\mathbf{ABX}) - \operatorname{tr}(\mathbf{X}^T\mathbf{B}^T\mathbf{A}^T\mathbf{X}) + \operatorname{tr}(\mathbf{X}^T\mathbf{B}^T\mathbf{A}^T\mathbf{ABX}) \\ &= \operatorname{tr}(\mathbf{X}^T\mathbf{X}) - 2\operatorname{tr}(\mathbf{X}^T\mathbf{ABX}) + \operatorname{tr}(\mathbf{X}^T\mathbf{B}^T\mathbf{A}^T\mathbf{ABX}) \end{aligned} \tag{119}$$

where we used that for any $\mathbf{G}, \mathbf{H} \in \mathbb{R}^{N\times N}$ it is true that $\operatorname{tr}(\mathbf{G}+\mathbf{H}) = \operatorname{tr}(\mathbf{G}) + \operatorname{tr}(\mathbf{H})$ and $\operatorname{tr}(\mathbf{G}^T) = \operatorname{tr}(\mathbf{G})$

Next we will use Equation 101 from Petersen and Pedersen (2012) which states that for any matrices $\mathbf{G}, \mathbf{H}, \mathbf{J} \in \mathbb{R}^{N\times N}$ we have

$$\frac{\partial}{\partial \mathbf{H}} \operatorname{tr}(\mathbf{GHJ}) = \mathbf{G}^T\mathbf{J}^T \tag{120}$$

and Equation 116 which states that for any matrices $\mathbf{G}, \mathbf{H}, \mathbf{J} \in \mathbb{R}^{N\times N}$ we have

$$\frac{\partial}{\partial H} \operatorname{tr}(\mathbf{G}^T\mathbf{H}^T\mathbf{JHG}) = \mathbf{J}^T\mathbf{HGG}^T + \mathbf{JHGG}^T \tag{121}$$

So computing the gradient of the RSS with respect to $\mathbf{A}$ we have

$$\begin{aligned} \mathbf{G}^{(\mathbf{A})} &= \nabla_{\mathbf{A}} \text{RSS} \\ &= \nabla_{\mathbf{A}} \left[\operatorname{tr}(\mathbf{X}^T\mathbf{X}) - 2\operatorname{tr}(\mathbf{X}^T\mathbf{ABX}) + \operatorname{tr}(\mathbf{X}^T\mathbf{B}^T\mathbf{A}^T\mathbf{ABX})\right] \\ &= -2\nabla_{\mathbf{A}} \operatorname{tr}(\underbrace{\mathbf{X}^T}_{G}\underbrace{\mathbf{A}}_{H}\underbrace{\mathbf{BX}}_{J}) + \nabla_{\mathbf{A}} \operatorname{tr}(\underbrace{(\mathbf{B}X)^T}_{\mathbf{G}^T}\underbrace{\mathbf{A}^T}_{H^T}\underbrace{\mathbf{I}}_{J}\underbrace{\mathbf{A}}_{H}\underbrace{\mathbf{BX}}_{G}) \\ &= -2\mathbf{XX}^T\mathbf{B}^T + \left(\mathbf{I}^T\mathbf{ABXX}^T\mathbf{B}^T + \mathbf{IABXX}^T\mathbf{B}^T\right) \\ &= -2\mathbf{XX}^T\mathbf{B}^T + 2\mathbf{ABXX}^T\mathbf{B}^T \\ &= 2\left(\mathbf{ABXX}^T\mathbf{B}^T - \mathbf{XX}^T\mathbf{B}^T\right) \\ &= 2\left(\mathbf{AZZ}^T - \mathbf{XZ}^T\right) \end{aligned} \tag{122}$$

Similarly, computing the gradient of the RSS with respect to $\mathbf{B}$ we have

$$\begin{aligned}
\mathbf{G}^{(\mathbf{B})} &= \nabla_{\mathbf{B}}\,\mathrm{RSS} \\
&= \nabla_{\mathbf{B}}\left[\mathrm{tr}(\mathbf{X}^T\mathbf{X}) - 2\,\mathrm{tr}(\mathbf{X}^T\mathbf{A}\mathbf{B}X) + \mathrm{tr}(\mathbf{X}^T\mathbf{B}^T\mathbf{A}^T\mathbf{A}\mathbf{B}\mathbf{X})\right] \\
&= -2\nabla_{\mathbf{B}}\,\mathrm{tr}(\underbrace{\mathbf{X}^T\mathbf{A}}_{G}\underbrace{\mathbf{B}}_{H}\underbrace{\mathbf{X}}_{J}) + \nabla_{\mathbf{B}}\,\mathrm{tr}(\underbrace{\mathbf{X}^T}_{\mathbf{G}^T}\underbrace{\mathbf{B}^T}_{H^T}\underbrace{\mathbf{A}^T\mathbf{A}}_{J}\underbrace{\mathbf{B}}_{H}\underbrace{\mathbf{X}}_{G}) \\
&= -2\mathbf{A}^T\mathbf{X}\mathbf{X}^T + \left(\mathbf{A}^T\mathbf{A}\mathbf{B}\mathbf{X}\mathbf{X}^T + \mathbf{A}^T\mathbf{A}\mathbf{B}\mathbf{X}\mathbf{X}^T\right) \\
&= -2\mathbf{A}^T\mathbf{X}\mathbf{X}^T + 2\mathbf{A}^T\mathbf{A}\mathbf{B}\mathbf{X}\mathbf{X}^T \\
&= 2\left(\mathbf{A}^T\mathbf{A}\mathbf{B}\mathbf{X}\mathbf{X}^T - \mathbf{A}^T\mathbf{X}\mathbf{X}^T\right)
\end{aligned} \tag{123}$$

#### 8.8 Gradient of l1-Normalization

Let $\mathbf{p} \in \mathbb{R}^K$. Define $\mathbf{r} \in \mathbb{R}^K$ element-wise as

$$r_k = \frac{\max(p_k, 0)}{\sum_{k''=1}^{K}\max(p_{k''}, 0)} = \frac{q_k}{\sum_{k''=1}^{K} q_{k''}} \tag{124}$$

where we define $q_k = \max(p_k, 0)$.

Then the gradient $\frac{d\mathbf{r}}{d\mathbf{p}} \in \mathbb{R}^{K\times K}$ is given by the chain rule

$$\frac{d\mathbf{r}}{d\mathbf{p}} = \frac{d\mathbf{r}}{d\mathbf{q}}\frac{d\mathbf{q}}{d\mathbf{p}} \tag{125}$$

The gradient of the rectified linear unit is given by

$$\frac{d\mathbf{q}}{d\mathbf{p}} = \mathrm{diag}\left(\begin{bmatrix}\mathbb{1}\left[p_1 \geq 0\right] & \dots & \mathbb{1}\left[p_K \geq 0\right]\end{bmatrix}\right) \tag{126}$$

While the gradient of the normalization is given by (see derivation below)

$$\frac{d\mathbf{r}}{d\mathbf{q}} = \frac{\left(\sum_{k''=1}^{K} q_{k''}\right) I_K - \mathbf{q}\mathbf{1}_K^T}{\left(\sum_{k''=1}^{K} q_{k''}\right)^2} \tag{127}$$

Thus we have

$$\frac{d\mathbf{r}}{d\mathbf{p}} = \frac{\left(\sum_{k''=1}^{K} q_{k''}\right) I_K - \mathbf{q}\mathbf{1}_K^T}{\left(\sum_{k''=1}^{K} q_{k''}\right)^2}\,\mathrm{diag}\begin{bmatrix}\mathbb{1}\left[p_1 \geq 0\right] & \dots & \mathbb{1}\left[p_K \geq 0\right]\end{bmatrix}^T \tag{128}$$

The gradient $\frac{d\mathbf{r}}{d\mathbf{p}} \in \mathbb{R}^{K\times K}$ has the form

$$\frac{d\mathbf{r}}{d\mathbf{q}} = \begin{bmatrix} \frac{\partial r_1}{\partial q_1} & \dots & \frac{\partial r_1}{\partial q_K} \\ \vdots & \ddots & \vdots \\ \frac{\partial r_K}{\partial q_1} & \dots & \frac{\partial r_K}{\partial q_K} \end{bmatrix} \tag{129}$$

The off-diagonal terms (i.e. $k \neq k'$) are given by

$$\begin{aligned}
\frac{\partial r_k}{\partial q_{k'}} &= \frac{\partial}{\partial q_{k'}} \frac{q_k}{\sum_{k''=1}^{K} q_{k''}} \\
&= q_k \frac{\partial}{\partial q_{k'}} \left( \sum_{k''=1}^{K} q_{k''} \right)^{-1} \\
&= q_k (-1) \left( \sum_{k''=1}^{K} q_{k''} \right)^{-2} \underbrace{\frac{\partial}{\partial q_{k'}} \left( \sum_{k''=1}^{K} q_{k''} \right)}_{=1} \\
&= -\frac{q_k}{\left( \sum_{k''=1}^{K} q_{k''} \right)^2}
\end{aligned} \tag{130}$$

To compute the diagonal terms we need to use the chain rule and can re-use the result from above

$$\begin{aligned}
\frac{\partial r_k}{\partial q_k} &= \frac{\partial}{\partial q_k} \frac{q_k}{\sum_{k''=1}^{K} q_{k''}} \\
&= \frac{\partial}{\partial q_k} \left[ \left( \frac{\partial}{\partial q_k} q_k \right) \left( \sum_{k''=1}^{K} q_{k''} \right)^{-1} \right] + \left[ q_k \left( \frac{\partial}{\partial q_k} \left( \sum_{k''=1}^{K} q_{k''} \right)^{-1} \right) \right] \\
&= \left[ 1 \left( \sum_{k''=1}^{K} q_{k''} \right)^{-1} \right] + \left[ -\frac{q_k}{\left( \sum_{k''=1}^{K} q_{k''} \right)^2} \right] \\
&= \frac{\sum_{k''=1}^{K} q_{k''}}{\left( \sum_{k''=1}^{K} q_{k''} \right)^2} - \frac{q_k}{\left( \sum_{k''=1}^{K} q_{k''} \right)^2}
\end{aligned} \tag{131}$$

Now putting everything together we have

$$\frac{d\mathbf{r}}{d\mathbf{q}} = \frac{\left( \sum_{k''=1}^{K} q_{k''} \right) I_K - \mathbf{q}\mathbf{1}_K^T}{\left( \sum_{k''=1}^{K} q_{k''} \right)^2} \tag{132}$$

#### 8.9 Gradient of Objective with Relaxed Archetype Constraints

Given the objective

$$\begin{aligned}
\hat{\mathbf{A}}, \hat{\mathbf{B}} = &\underset{\substack{\mathbf{A} \in \mathbb{R}^{N \times K} \\ \mathbf{B} \in \mathbb{R}^{K \times N}}}{\arg\min} \|\mathbf{X} - \mathbf{A} \operatorname{diag}(\boldsymbol{\alpha}) \mathbf{B} \mathbf{X}\|_F^2 \quad \text{subject to} \\
&\mathbf{A} \geq 0, \quad \mathbf{A}\mathbf{1}_K = \mathbf{1}_N \\
&\mathbf{B} \geq 0, \quad \mathbf{B}\mathbf{1}_N = \mathbf{1}_K \\
&\forall k \in \{1, ..., K\} \quad 1 - \delta \leq \alpha_k \leq 1 + \delta
\end{aligned} \tag{133}$$

we seek to derive the gradient with respect to $\mathbf{A}$, $\mathbf{B}$, and $\boldsymbol{\alpha}$. Following section 8.7 we first rewrite the objective in terms of the trace.

$$\begin{aligned}\text{RSS} &= \|\mathbf{X} - \mathbf{A}\operatorname{diag}(\boldsymbol{\alpha})\mathbf{B}\mathbf{X}\|_F^2 \\ &= \operatorname{tr}\left((\mathbf{X} - \mathbf{A}\operatorname{diag}(\boldsymbol{\alpha})\mathbf{B}\mathbf{X})^T(\mathbf{X} - \mathbf{A}\operatorname{diag}(\boldsymbol{\alpha})\mathbf{B}\mathbf{X})\right) \\ &= \operatorname{tr}\left(\mathbf{X}^T\mathbf{X}\right) - \operatorname{tr}\left(\mathbf{X}^T\mathbf{A}\operatorname{diag}(\boldsymbol{\alpha})\mathbf{B}\mathbf{X}\right) - \operatorname{tr}\left(\mathbf{X}^T\mathbf{B}^T\operatorname{diag}(\boldsymbol{\alpha})\mathbf{A}^T\mathbf{X}\right) \\ &\quad + \operatorname{tr}\left(\mathbf{X}^T\mathbf{B}^T\operatorname{diag}(\boldsymbol{\alpha})\mathbf{A}^T\mathbf{A}\operatorname{diag}(\boldsymbol{\alpha})\mathbf{B}\mathbf{X}\right) \\ &= \operatorname{tr}\left(\mathbf{X}^T\mathbf{X}\right) - 2\operatorname{tr}\left(\mathbf{X}^T\mathbf{A}\operatorname{diag}(\boldsymbol{\alpha})\mathbf{B}\mathbf{X}\right) + \operatorname{tr}\left(\mathbf{X}^T\mathbf{B}^T\operatorname{diag}(\boldsymbol{\alpha})\mathbf{A}^T\mathbf{A}\operatorname{diag}(\boldsymbol{\alpha})\mathbf{B}\mathbf{X}\right)\end{aligned} \tag{134}$$

We will use again the identities from Equation 120 and Equation 121

Computing the gradient of the RSS with respect to $\mathbf{A}$ we have

$$\begin{aligned}\mathbf{G}^{(A)} &= \nabla_{\mathbf{A}}\,\text{RSS} \\ &= \nabla_{\mathbf{A}}\left[\operatorname{tr}\left(\mathbf{X}^T\mathbf{X}\right) - 2\operatorname{tr}\left(\mathbf{X}^T\mathbf{A}\operatorname{diag}(\boldsymbol{\alpha})\mathbf{B}\mathbf{X}\right) + \operatorname{tr}\left(\mathbf{X}^T\mathbf{B}^T\operatorname{diag}(\boldsymbol{\alpha})\mathbf{A}^T\mathbf{A}\operatorname{diag}(\boldsymbol{\alpha})\mathbf{B}\mathbf{X}\right)\right] \\ &= -2\nabla_{\mathbf{A}}\operatorname{tr}(\underbrace{\mathbf{X}^T}_{G}\underbrace{\mathbf{A}}_{H}\underbrace{\operatorname{diag}(\boldsymbol{\alpha})\mathbf{B}\mathbf{X}}_{J}) + \nabla_{\mathbf{A}}\operatorname{tr}(\underbrace{(\operatorname{diag}(\boldsymbol{\alpha})\mathbf{B}\mathbf{X})^T}_{\mathbf{G}^T}\underbrace{\mathbf{A}^T}_{H^T}\underbrace{\mathbf{I}}_{J}\underbrace{\mathbf{A}}_{H}\underbrace{\operatorname{diag}(\boldsymbol{\alpha})\mathbf{B}\mathbf{X}}_{G}) \\ &= -2\mathbf{X}\mathbf{X}^T\mathbf{B}^T\operatorname{diag}(\boldsymbol{\alpha}) + 2\mathbf{A}\operatorname{diag}(\boldsymbol{\alpha})\mathbf{B}\mathbf{X}\mathbf{X}^T\mathbf{B}^T\operatorname{diag}(\boldsymbol{\alpha}) \\ &= 2\left(\mathbf{A}\operatorname{diag}(\boldsymbol{\alpha})\mathbf{B}\mathbf{X}\mathbf{X}^T\mathbf{B}^T\operatorname{diag}(\boldsymbol{\alpha}) - \mathbf{X}\mathbf{X}^T\mathbf{B}^T\operatorname{diag}(\boldsymbol{\alpha})\right)\end{aligned} \tag{135}$$

Computing the gradient of the RSS with respect to $\mathbf{B}$ we have

$$\begin{aligned}\mathbf{G}^{(B)} &= \nabla_{\mathbf{B}}\,\text{RSS} \\ &= \nabla_{\mathbf{B}}\left[\operatorname{tr}\left(\mathbf{X}^T\mathbf{X}\right) - 2\operatorname{tr}\left(\mathbf{X}^T\mathbf{A}\operatorname{diag}(\boldsymbol{\alpha})\mathbf{B}\mathbf{X}\right) + \operatorname{tr}\left(\mathbf{X}^T\mathbf{B}^T\operatorname{diag}(\boldsymbol{\alpha})\mathbf{A}^T\mathbf{A}\operatorname{diag}(\boldsymbol{\alpha})\mathbf{B}\mathbf{X}\right)\right] \\ &= -2\nabla_{\mathbf{B}}\operatorname{tr}(\underbrace{\mathbf{X}^T\mathbf{A}\operatorname{diag}(\boldsymbol{\alpha})}_{G}\underbrace{\mathbf{B}}_{H}\underbrace{\mathbf{X}}_{J}) + \nabla_{\mathbf{B}}\operatorname{tr}(\underbrace{\mathbf{X}^T}_{\mathbf{G}^T}\underbrace{\mathbf{B}^T}_{H^T}\underbrace{\operatorname{diag}(\boldsymbol{\alpha})\mathbf{A}^T\mathbf{A}\operatorname{diag}(\boldsymbol{\alpha})}_{J}\underbrace{\mathbf{B}}_{H}\underbrace{\mathbf{X}}_{G}) \\ &= -2\operatorname{diag}(\boldsymbol{\alpha})\mathbf{A}^T\mathbf{X}\mathbf{X}^T + 2\operatorname{diag}(\boldsymbol{\alpha})\mathbf{A}^T\mathbf{A}\operatorname{diag}(\boldsymbol{\alpha})\mathbf{B}\mathbf{X}\mathbf{X}^T \\ &= 2\left[\operatorname{diag}(\boldsymbol{\alpha})\mathbf{A}^T\mathbf{A}\operatorname{diag}(\boldsymbol{\alpha})\mathbf{B}\mathbf{X}\mathbf{X}^T - \operatorname{diag}(\boldsymbol{\alpha})\mathbf{A}^T\mathbf{X}\mathbf{X}^T\right]\end{aligned} \tag{136}$$

Finally, computing the gradient of the RSS with respect to $\boldsymbol{\alpha}$ we have

$$\begin{aligned}\mathbf{G}^{(\alpha)} &= \nabla_{\alpha}\,\text{RSS} \\ &= \nabla_{\alpha}\left[\operatorname{tr}\left(\mathbf{X}^T\mathbf{X}\right) - 2\operatorname{tr}\left(\mathbf{X}^T\mathbf{A}\operatorname{diag}(\boldsymbol{\alpha})\mathbf{B}\mathbf{X}\right) + \operatorname{tr}\left(\mathbf{X}^T\mathbf{B}^T\operatorname{diag}(\boldsymbol{\alpha})\mathbf{A}^T\mathbf{A}\operatorname{diag}(\boldsymbol{\alpha})\mathbf{B}\mathbf{X}\right)\right] \\ &= -2\nabla_{\alpha}\operatorname{tr}(\underbrace{\mathbf{X}^T\mathbf{A}}_{G}\underbrace{\operatorname{diag}(\boldsymbol{\alpha})}_{H}\underbrace{\mathbf{B}\mathbf{X}}_{J}) + \nabla_{\alpha}\operatorname{tr}(\underbrace{\mathbf{X}^T\mathbf{B}^T}_{\mathbf{G}^T}\underbrace{\operatorname{diag}(\boldsymbol{\alpha})}_{H^T}\underbrace{\mathbf{A}^T\mathbf{A}}_{J}\underbrace{\operatorname{diag}(\boldsymbol{\alpha})}_{H}\underbrace{\mathbf{B}\mathbf{X}}_{G}) \\ &= -2\operatorname{diag}\left(\mathbf{A}^T\mathbf{X}\mathbf{X}^T\mathbf{B}^T\right) + 2\operatorname{diag}\left(\mathbf{A}^T\mathbf{A}\operatorname{diag}(\boldsymbol{\alpha})\mathbf{B}\mathbf{X}\mathbf{X}^T\mathbf{B}^T\right) \\ &= 2\operatorname{diag}\left(\mathbf{A}^T\mathbf{A}\operatorname{diag}(\boldsymbol{\alpha})\mathbf{B}\mathbf{X}\mathbf{X}^T\mathbf{B}^T - \mathbf{A}^T\mathbf{X}\mathbf{X}^T\mathbf{B}^T\right)\end{aligned} \tag{137}$$

#### 8.10 Gradient of Objective with Relaxed Archetype Constraints and Coresets

Given the objective

$$\begin{aligned}&\hat{\mathbf{A}}, \hat{\mathbf{B}} = \arg\min \|\mathbf{W}\tilde{\mathbf{X}} - \mathbf{W}\mathbf{A}\operatorname{diag}(\boldsymbol{\alpha})\mathbf{B}\tilde{\mathbf{X}}\|_F^2 \quad \text{subject to} \\ &\mathbf{A} \in F(\tilde{N}, K) \\ &\mathbf{B} \in F(K, \tilde{N}) \\ &\forall k \in \{1, ..., K\} \quad 1 - \delta \leq \alpha_k \leq 1 + \delta\end{aligned} \tag{138}$$

we seek to derive the gradient with respect to $\mathbf{A}$, $\mathbf{B}$, and $\boldsymbol{\alpha}$.

To simplify the derivation we define $\breve{\mathbf{A}} = \mathbf{W}\mathbf{A}$ and $\breve{\mathbf{X}} = \mathbf{W}\tilde{\mathbf{X}}$. Following section 8.7 we first rewrite the objective in terms of the trace.

$$\begin{aligned}
\text{RSS} &= \|\breve{\mathbf{X}} - \breve{\mathbf{A}}\,\text{diag}(\boldsymbol{\alpha})\mathbf{B}\tilde{\mathbf{X}}\|_F^2 \\
&= \text{tr}\left(\left(\breve{\mathbf{X}} - \breve{\mathbf{A}}\,\text{diag}(\boldsymbol{\alpha})\mathbf{B}\tilde{\mathbf{X}}\right)^T\left(\breve{\mathbf{X}} - \breve{\mathbf{A}}\,\text{diag}(\boldsymbol{\alpha})\mathbf{B}\tilde{\mathbf{X}}\right)\right) \\
&= \text{tr}\left(\breve{\mathbf{X}}^T\breve{\mathbf{X}}\right) - \text{tr}\left(\breve{\mathbf{X}}^T\breve{\mathbf{A}}\,\text{diag}(\boldsymbol{\alpha})\mathbf{B}\tilde{\mathbf{X}}\right) - \text{tr}\left(\tilde{\mathbf{X}}^T\mathbf{B}^T\,\text{diag}(\boldsymbol{\alpha})\breve{\mathbf{A}}^T\breve{\mathbf{X}}\right) \\
&\quad + \text{tr}\left(\tilde{\mathbf{X}}^T\mathbf{B}^T\,\text{diag}(\boldsymbol{\alpha})\breve{\mathbf{A}}^T\breve{\mathbf{A}}\,\text{diag}(\boldsymbol{\alpha})\mathbf{B}\tilde{\mathbf{X}}\right) \\
&= \text{tr}\left(\breve{\mathbf{X}}^T\breve{\mathbf{X}}\right) - 2\,\text{tr}\left(\breve{\mathbf{X}}^T\breve{\mathbf{A}}\,\text{diag}(\boldsymbol{\alpha})\mathbf{B}\tilde{\mathbf{X}}\right) + \text{tr}\left(\tilde{\mathbf{X}}^T\mathbf{B}^T\,\text{diag}(\boldsymbol{\alpha})\breve{\mathbf{A}}^T\breve{\mathbf{A}}\,\text{diag}(\boldsymbol{\alpha})\mathbf{B}\tilde{\mathbf{X}}\right)
\end{aligned} \tag{139}$$

Using the identities from Equation 120 and Equation 121, the gradient of the RSS with respect to $\mathbf{B}$ is given by

$$\begin{aligned}
\mathbf{G}^{(B)} &= \nabla_{\mathbf{B}}\,\text{RSS} \\
&= \nabla_{\mathbf{B}}\left[\text{tr}\left(\breve{\mathbf{X}}^T\breve{\mathbf{X}}\right) - 2\,\text{tr}\left(\breve{\mathbf{X}}^T\breve{\mathbf{A}}\,\text{diag}(\boldsymbol{\alpha})\mathbf{B}\tilde{\mathbf{X}}\right) + \text{tr}\left(\tilde{\mathbf{X}}^T\mathbf{B}^T\,\text{diag}(\boldsymbol{\alpha})\breve{\mathbf{A}}^T\breve{\mathbf{A}}\,\text{diag}(\boldsymbol{\alpha})\mathbf{B}\tilde{\mathbf{X}}\right)\right] \\
&= -2\nabla_{\mathbf{B}}\,\text{tr}(\underbrace{\breve{\mathbf{X}}^T\breve{\mathbf{A}}\,\text{diag}(\boldsymbol{\alpha})}_{G}\underbrace{\mathbf{B}}_{H}\underbrace{\tilde{\mathbf{X}}}_{J}) + \nabla_{\mathbf{B}}\,\text{tr}(\underbrace{\tilde{\mathbf{X}}^T}_{\mathbf{G}^T}\underbrace{\mathbf{B}^T}_{H^T}\underbrace{\text{diag}(\boldsymbol{\alpha})\breve{\mathbf{A}}^T\breve{\mathbf{A}}\,\text{diag}(\boldsymbol{\alpha})}_{J}\underbrace{\mathbf{B}}_{H}\underbrace{\tilde{\mathbf{X}}}_{G}) \\
&= -2\,\text{diag}(\boldsymbol{\alpha})\breve{\mathbf{A}}^T\breve{\mathbf{X}}\tilde{\mathbf{X}}^T + 2\,\text{diag}(\boldsymbol{\alpha})\breve{\mathbf{A}}^T\breve{\mathbf{A}}\,\text{diag}(\boldsymbol{\alpha})\mathbf{B}\tilde{\mathbf{X}}\tilde{\mathbf{X}}^T \\
&= 2\left[\text{diag}(\boldsymbol{\alpha})\breve{\mathbf{A}}^T\breve{\mathbf{A}}\,\text{diag}(\boldsymbol{\alpha})\mathbf{B}\tilde{\mathbf{X}}\tilde{\mathbf{X}}^T - \text{diag}(\boldsymbol{\alpha})\breve{\mathbf{A}}^T\breve{\mathbf{X}}\tilde{\mathbf{X}}^T\right]
\end{aligned} \tag{140}$$

Computing the gradient of the RSS with respect to $\boldsymbol{\alpha}$ we have

$$\begin{aligned}
\mathbf{G}^{(\alpha)} &= \nabla_{\boldsymbol{\alpha}}\,\text{RSS} \\
&= \nabla_{\boldsymbol{\alpha}}\left[\text{tr}\left(\breve{\mathbf{X}}^T\breve{\mathbf{X}}\right) - 2\,\text{tr}\left(\breve{\mathbf{X}}^T\breve{\mathbf{A}}\,\text{diag}(\boldsymbol{\alpha})\mathbf{B}\tilde{\mathbf{X}}\right) + \text{tr}\left(\tilde{\mathbf{X}}^T\mathbf{B}^T\,\text{diag}(\boldsymbol{\alpha})\breve{\mathbf{A}}^T\breve{\mathbf{A}}\,\text{diag}(\boldsymbol{\alpha})\mathbf{B}\tilde{\mathbf{X}}\right)\right] \\
&= -2\nabla_{\boldsymbol{\alpha}}\,\text{tr}(\underbrace{\breve{\mathbf{X}}^T\breve{\mathbf{A}}}_{G}\underbrace{\text{diag}(\boldsymbol{\alpha})}_{H}\underbrace{\mathbf{B}\tilde{\mathbf{X}}}_{J}) + \nabla_{\boldsymbol{\alpha}}\,\text{tr}(\underbrace{\tilde{\mathbf{X}}^T\mathbf{B}^T}_{\mathbf{G}^T}\underbrace{\text{diag}(\boldsymbol{\alpha})}_{H^T}\underbrace{\breve{\mathbf{A}}^T\breve{\mathbf{A}}}_{J}\underbrace{\text{diag}(\boldsymbol{\alpha})}_{H}\underbrace{\mathbf{B}\tilde{\mathbf{X}}}_{G}) \\
&= -2\,\text{diag}\left(\breve{\mathbf{A}}^T\breve{\mathbf{X}}\tilde{\mathbf{X}}^T\mathbf{B}^T\right) + 2\,\text{diag}\left(\breve{\mathbf{A}}^T\breve{\mathbf{A}}\,\text{diag}(\boldsymbol{\alpha})\mathbf{B}\tilde{\mathbf{X}}\tilde{\mathbf{X}}^T\mathbf{B}^T\right) \\
&= 2\,\text{diag}\left(\breve{\mathbf{A}}^T\breve{\mathbf{A}}\,\text{diag}(\boldsymbol{\alpha})\mathbf{B}\tilde{\mathbf{X}}\tilde{\mathbf{X}}^T\mathbf{B}^T - \breve{\mathbf{A}}^T\breve{\mathbf{X}}\tilde{\mathbf{X}}^T\mathbf{B}^T\right)
\end{aligned} \tag{141}$$
